# Hybrid dynamic pharmacophore models as effective tools to identify novel chemotypes for anti-TB inhibitor design: A case study with Mtb-DapB

**DOI:** 10.1101/2020.01.20.912063

**Authors:** Chinmayee Choudhury, Anshu Bhardwaj

## Abstract

Antimicrobial resistance (AMR) is one of the most serious global public health threats as it compromises the successful treatment of deadly infectious diseases like tuberculosis. New therapeutics are constantly needed but it takes a long time and is expensive to explore new biochemical space. One way to address this issue is to repurpose the validated targets and identify novel chemotypes that can simultaneously bind to multiple binding pockets of these targets as a new lead generation strategy. This study reports such a strategy, dynamic hybrid pharmacophore model (DHPM), which represents the combined interaction features of different binding pockets contrary to the conventional approaches, where pharmacophore models are generated from single binding sites. We have considered Mtb-DapB, a validated mycobacterial drug target, as our model system to explore the effectiveness of DHPMs to screen novel unexplored compounds. Mtb-DapB has a cofactor binding site (CBS) and an adjacent substrate binding site (SBS). Four different model systems of Mtb-DapB were designed where, either NADPH/NADH occupies CBS in presence/absence of an inhibitor 2, 6-PDC in the adjacent SBS. Two more model systems were designed, where 2, 6-PDC was linked to NADPH and NADH to form hybrid molecules. The six model systems were subjected to 200ns molecular dynamics simulations and trajectories were analyzed to identify stable ligand-receptor interaction features. Based on these interactions, conventional pharmacophore models (CPM) were generated from the individual binding sites while DHPMs were created from hybrid-molecules occupying both binding sites. A huge library of 15, 63,764 publicly available molecules were screened by CPMs and DHPMs. The screened hits obtained from both types of models were compared based on their Hashed binary molecular fingerprints and 4-point pharmacophore fingerprints using Tanimoto, Cosine, Dice and Tversky similarity matrices. Molecules screened by DHPM exhibited significant structural diversity, better binding strength and drug like properties as compared to the compounds screened by CPMs indicating the efficiency of DHPM to explore new chemical space for anti-TB drug discovery. The idea of DHPM can be applied for a wide range of mycobacterial or other pathogen targets to venture into unexplored chemical space.

## 1 Introduction

Tuberculosis (TB) is the leading cause of death worldwide due to a single infectious agent (Global Tuberculosis Report 2019),(Balganesh et al., 2008). The standard therapies of treating TB with a combination of several antibiotics over a period of six to nine months (Dutt and Stead, 1997),(Zumla et al., 2015) and have severe limitations due to compliance which leads to emergence of drug resistant *Mycobacterium tuberculosis* (Mtb) (Prasad and Srivastava, 2013). Antimicrobial resistance (AMR) is one of the most serious global public health threats (Passi et al., 2019). The “End Tuberculosis Strategy” (Global Tuberculosis Report 2019) of WHO calls for intensified research and innovation in TB drug discovery using multidisciplinary approaches to identify novel drug targets as well as fast and accurate techniques to design new chemical entities with higher potency to address the scourge of Mtb. In the last few decades, there has been rapid development in computational drug discovery algorithms for *ab initio* protein modeling, homology modeling, protein folding dynamics, molecular docking, pharmacophore modeling, virtual screening, quantitative structure activity relationship (QSAR) etc. (Choudhury et al., 2014) (Choudhury et al., 2015) (Choudhury et al., 2016) (Choudhury et al., 2016) (Gaur et al., 2017) (Kumar Srivastava et al., 2012) (Kurumurthy et al., 2012). Several new strategies are also designed for repurposing existing drugs or discover new ones (Passi et al., 2018). The increasing number of experimental structures of drug targets deposited in the protein data bank (PDB) and atomistic details of the binding modes of co-crystalized compounds/drugs/ natural ligands (substrates/cofactors) provide a strong primer for structure-based lead generation and optimization. However, choosing the right molecular interaction features in the active sites of a drug target is the most important step of structure-based drug design (Wieder et al., 2017). Most often, the interactions made by the natural substrate(s) of the target protein/enzymes as reported in their static experimental structures are considered for designing new inhibitors. However, for target inhibition by competitive binding via specific interactions in the binding pocket, the static structure does not provide information on all potential and stable molecular interactions in the binding site. Also, this approach offers limited specificity and chemical diversity when cofactor/substrates are involved as these are common to both the host and the pathogen. Hence, utilization of additional information on adjacent binding pockets and their ligand binding potential along with the dynamics of the binding pockets might prove beneficial to design/screen novel chemotypes. These chemotypes can form specific stable interactions with multiple binding sites of the target offering better specificity and scope for introducing chemical diversity (Duckworth et al., 2011 & San Jose et al., 2013). In recent years, target specific interactions in the form of pharmacophore models for virtual screening gained immense popularity (Schuetz et al., 2018) (Schaller et al., 2020).

In this study we have considered DapB, one of the validated drug targets in Mtb. DapB enzyme belongs to Diaminopimelate (DAP) biosynthetic pathway (Cox, 1996) (Cox et al., 2000) (Vashisht et al., 2012) and inhibition of this enzyme blocks the production of meso-diaminopimelate thus leading to inhibition of *de novo* lysine biosynthesis and peptidoglycan assembly (Figure 1(A)). Both of these pathways are crucial for the survival of the pathogen (Usha et al., 2012). Several groups made efforts to identify inhibitors of Mtb-DapB by exploring the potential of product analogues (Singh et al., 2013) as potential inhibitors, but have met with very limited success. Pyridine-2, 6-dicaboxylate (2, 6-PDC) and few other heterocyclic aromatic product analogues have been identified with IC50 as high as 26 µM for Mtb-DapB. Also, a number of sulphonamide inhibitors of Mtb-DapB (substrate analog for 2,3-dihydrodipicolinic acid) were also identified with Ki values ranging from 7 to 48 µM (Paiva et al., 2001). This indicates that only substrate or product mimicry is not sufficient and new lead generation strategies should be employed exploring the key features of the binding sites of the enzymes.

**Figure 1.**
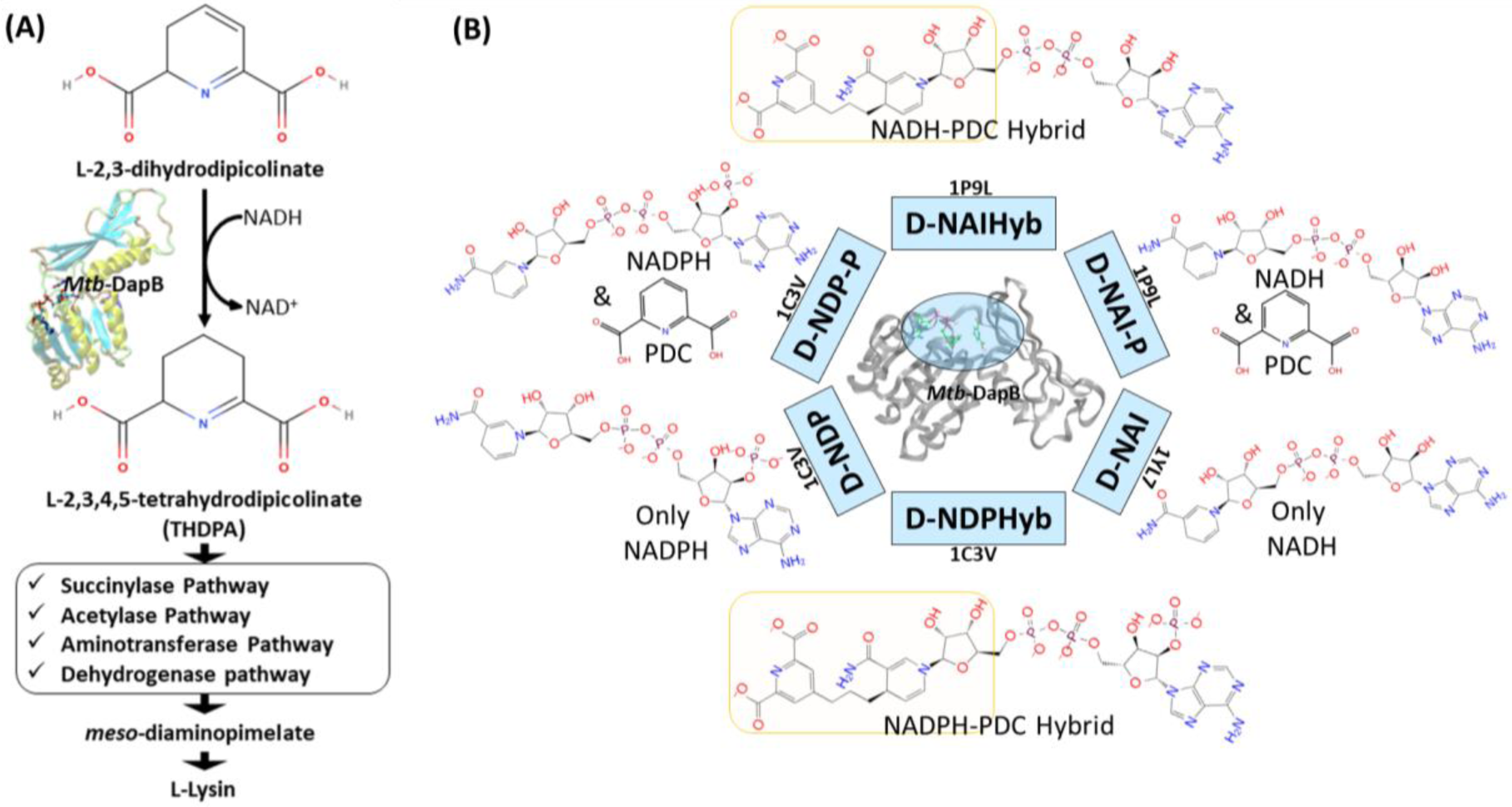
**(A)** Schematic diagram showing the role of Mtb-DapB in de novo lysine biosynthesis and peptidoglycan assembly, **(B)** Mtb-DapB model systems designed in the current study.

Molecular dynamics (MD) based pharmacophore models have emerged as quite powerful tools which not only account for the flexibility of the target(s), but also help to identify novel key interaction features in the binding sites (Spyrakis et al., 2015) which is otherwise unexplored in the crystal structures and may be harnessed to design new inhibitors (Guterres and Im, 2020), (Choudhury et al., 2014), (Choudhury et al., 2015) (Choudhury and Bhardwaj, 2020). Literature reports several studies that have used molecular dynamics to sample huge number of receptor conformations and generated pharmacophore models from these conformations which showed improved enrichment for specific interaction features (Wieder et al., 2016) (Culletta et al., 2020) (Perricone et al., 2017) (Tutone et al., 2019) (Bottegoni et al., 2011). MD simulations combined with virtual screening of compound libraries have led to identification of biologically active compounds for several targets (Tutone et al., 2016) (Tutone et al., 2014). In this study, hybrid pharmacophore models representing multiple ligand binding regions of Mtb-DapB were designed based on the stable molecular interactions identified from the multiple MD trajectories of the target bound to the co-crystal ligand molecules as well as hybrids of two such ligands bound to different binding sites (Figure 1(B)). We comparatively investigated the efficacies of the dynamics-based hybrid pharmacophore models (DHPM) based on multiple binding sites and the conventional pharmacophore models obtained from a single binding site, to obtain novel, target specific and diverse chemotypes with higher binding affinities as compared to the existing ones.

## 2 Material and Methods

### 2.1 Model Systems and Molecular Dynamics (MD) Simulations

Six different model systems were designed from three available experimental structures of Mtb-DapB for the study. The X-ray crystal structure 1C3V (Cirilli et al., 2003) was used to design three model systems, D-NDP-P, D-NDP and D-NDPHyb. The first one (D-NDP-P) is Mtb-DapB bound to NADPH (natural cofactor of Mtb-DapB) and 2,6-PDC (a mimetic of the natural substrate of Mtb-DapB) as in the reported structure, the second one (D-NDP) bound to only NADPH and the third one (D-NDPHyb) bound to a hybrid molecule created by linking NADPH and 2,6-PDC with a simple n-propyl linker. Similarly, the X-ray crystal structure 1P9L (Cirilli et al., 2003) was used to generate two model systems, D-NAI-P and D-NAIHyb. The first one (D-NAI-P) is Mtb-DapB bound to NADH (another natural ligand of Mtb-DapB) and 2,6-PDC as in the reported structure and the other model (D-NAIHyb) with a hybrid molecule created by linking NADH and 2,6-PDC with the n-propyl linker. The third crystal structure 1YL7 (Janowski et al., 2010) was used as in the reported structure bound to only NADH (D-NAI). Figure 1(B) depicts the model systems designed for this study.

200ns MD simulations were carried out under NPT ensemble on each model system using OPLS force field (Harder et al., 2016) with the Desmond MD simulation package (release 2017) (Shaw et al., 2014). Complete details of the MD Simulation methodology are given in Section S1 of the supplementary information.

### 2.2 MD Trajectory clustering and Generation of e-pharmacophore models

Hierarchical clustering was performed on each trajectory (RMSD 3Å) to sample representative structures with minimum RMSD of all the heavy atoms of the binding sites (residues within 5Å of the bound ligands). The protein ligand interactions were analyzed for each cluster representative and the ones with the most stable interaction features were considered further for pharmacophore modelling using the e-Pharmacophore option from the Phase (Dixon et al., 2006a) module of Schrodinger Suite. To generate energy-based e-Pharmacophore models (Salam et al., 2009) from the given protein ligand complexes, Phase first estimates the Glide extra precision (XP) (Friesner et al., 2006) energy terms for hydrophobic enclosure, hydrophobically packed correlated hydrogen bonds, electrostatic rewards, π-π stacking, cation-π, and other interactions. Each interaction is represented by a pharmacophore feature site and is assigned an energetic value equal to the sum of the Glide XP contributions of the atoms comprising the site. Then, the sites are ranked based on the energetic terms. Minimum inter-feature distance was maintained to be 2Å, while minimum inter-feature distances between the same types of features were assigned to be 4Å and the donors were presented as vectors. Receptor based excluded volume shells of 5Å thickness were created using the Van der Waals radii of the receptor atoms. The scaling factor was assigned as 0.50 and receptor atoms within 2Å of the ligand were ignored while defining the excluded volume shells as per the default settings. Conventional pharmacophore models were generated individually from NADPH, NADH and PDC from model systems D-NDP-P, D-NDP and D-NAI-P. Maximum number of features were assigned as 7 for these conventional models. None of the cluster representatives from the trajectory of D-NAI was used for pharmacophore model generation as none of the interactions formed between NADH and Mtb-DapB in this system were stable (occupancy < 40%). First type of hybrid pharmacophore models were generated directly from the hybrid molecules in model systems D-NDPHyb and D-NAIHyb. The maximum number of features were initially assigned as 10 for these hybrid models, but best 6 to 7 features were retained from the desired regions (highlighted in yellow in Figure 1(B)) of the hybrid molecules, after discarding the features associated with the adenosine and phosphate regions. The second type of hybrid pharmacophore models with 6-7 features were obtained by first merging the conventional models from NADPH/NADH and PDC and then discarding the features associated with ADP regions of NADPH/NADH. The pharmacophore models obtained individually from 2, 6-PDC were having only 3 to 4 features, which would not confer specificity to the models. So, they were not used further for ligand screening. The conventional pharmacophore models obtained from NADPH/NADH individually were named as N-type models while the two types of hybrid pharmacophore models were named as H-type pharmacophore models.

### 2.3 Pharmacophore screening

The ZINC natural product subsets with 132883 molecules (ZINC) and Asinex screening library (Asinex.com – Asinex Focused Libraries, Screening compounds, Pre-plated Sets) consisting of 530881 molecules was selected for our study. The Asinex library included the “BioDesign” library of 175851 pharmacologically relevant natural product like compounds, “lead-like” library with 91473 compounds (those have been screened against a panel of early ADMET tests including DMSO and water solubility, PAMPA, PGP and CYP inhibition) and “Gold and Platinum Collections” with 263557 molecules having a high degree of drug-likeness, in accordance with Lipinski’s Rule of 5. These molecules were subjected to preparation in LigPrep (Shelley et al., 2007), generating their ionization states at pH 7.0 (± 2.0) using Epik ionizer and five lowest energy conformers were retained for each compound. Then, these molecules were screened against the N-type and H-type pharmacophore models using the “Ligand and Database Screening” option of Phase module of Schrodinger Suite (Dixon et al., 2006a). 20 conformers were generated per molecule and the outputs were minimized before alignment against the pharmacophore models. The negative and acceptor features were considered equivalent and the minimum number of sites the molecule must match was assigned to be 5. Among many conformers of a ligand, the one with the best fitness score (S) evaluated by a specific fitness function (Dixon et al., 2006b) was retained for each compound. The consolidated lists of all the compounds screened by the N- and H-type models were named as N-set and H-set respectively. Various cheminformatics analyses were performed to compare the structural, physicochemical and ADMET properties of the N- and H-set. Mtb-DapB binding affinities of N- and H-set were estimated by molecular docking studies and compared.

### 2.4 Generation of molecular fingerprints and library comparison

Different types of hashed binary molecular fingerprints such as liner, radial, MOLPRINT2D and 4-point 3D pharmacophoric fingerprints were generated for N- and H-set molecules with Schrodinger. Then, the H-set molecules were compared with the N-set compounds on the basis of the above molecular finger prints using Tanimoto, Cosine, Dice and Tversky similarity matrices. Max Similarity (MaxSim) scores (calculated using Tanimoto, Cosine, Dice and Tversky similarity matrices) were obtained for each molecule of the query set (H-set) with respect to the N-set molecules. MaxSim represents the similarity score of each molecule from H-set with the nearest neighbor of the N-set. Various drug likeliness and ADMET properties were also calculated using QuikProp module of Schrodinger for the two sets of molecules and were compared to each other.

### 2.5 Molecular docking

The N- and H-set compounds were subjected to blind molecular docking with the cluster representatives used for pharmacophore model generation. As all the structures were prepared before MD simulations, these cluster representatives were directly used for grid generation. ‘Receptor Grid Generation’ module of Schrödinger was utilized to define interaction grids for molecular docking keeping the centroids of all residues within 5Å of both the bound ligands as grid centers. The size of the interaction grid, which covered almost the whole protein including both the binding sites was fixed to 20 Å for inner box and 24Å as outer box to facilitate a blind docking. Then, the N- and H-set compounds were subjected to docking calculations with the interaction grids of the selected cluster representatives using the Glide (Friesner et al., 2006) module of Schrödinger software package first with standard precision (SP) followed by eXtra Precision (XP) modes. Five best energy poses were generated for each compound. OPLS_2005 force field was used for docking with all default parameters. The resultant complexes were further submitted for binding energy estimation, where Molecular Mechanics-Generalized Born Surface Area (MM/GBSA) based binding free energy (ΔGbind) were computed for the complex using Prime module. The N- and H-type compounds were then compared based on their DapB binding affinities (Docking scores and ΔGbind) as well as their potential to make interactions with key residues as identified from the MD trajectories. Figure 2 shows the overall workflow of the study.

**Figure 2.**
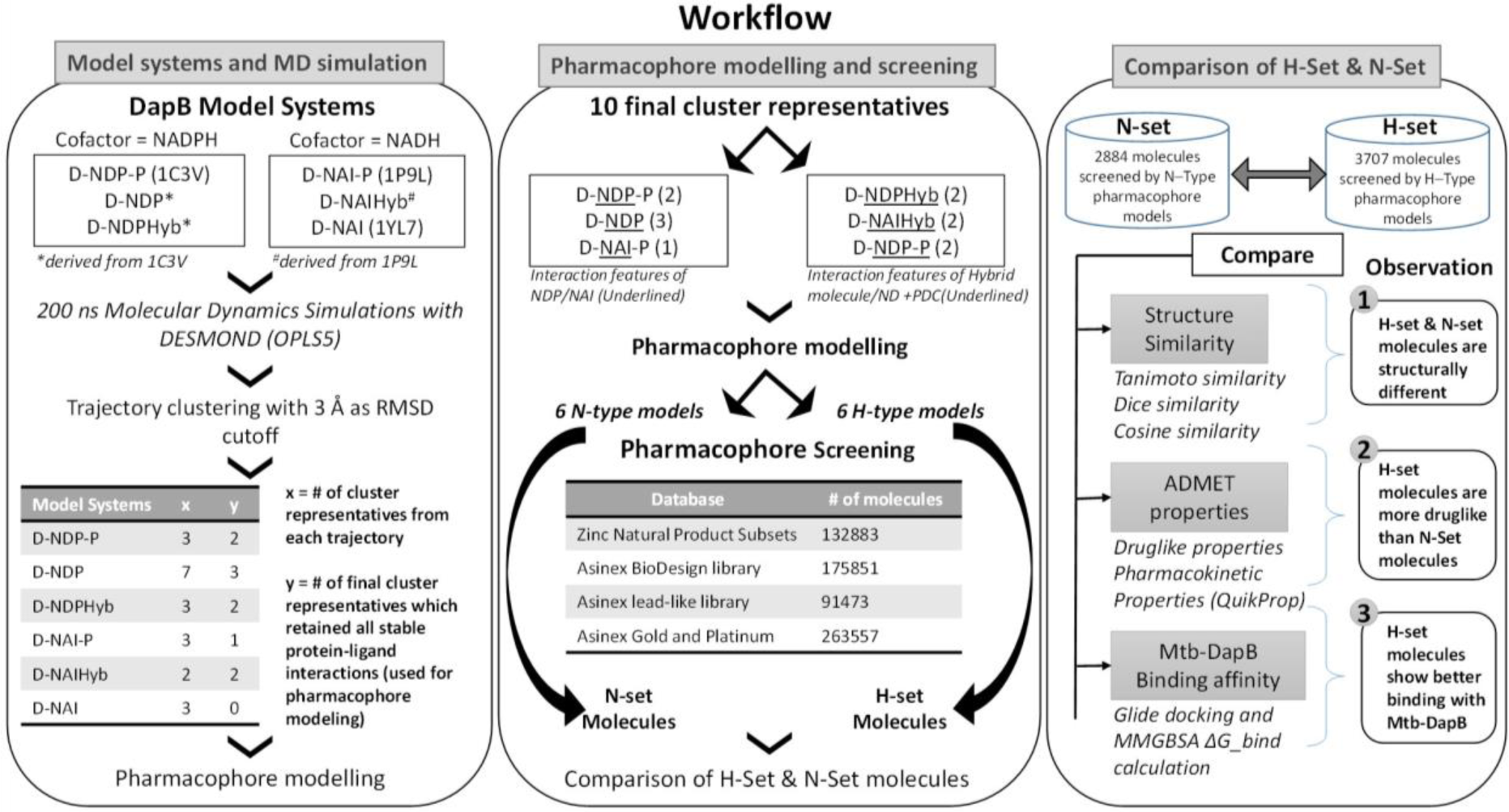
Overall workflow of the study.

## 3 Results and Discussion

### 3.1 Overall Structure of Mtb-DapB model systems

The Mtb-DapB is a homo-tetramer of 245-residue monomers (Fig 3(A)) with two major domains connected with two short hinge regions. The N-terminal domain (1-106 and 216-245) contains six β-strands (β1-β5 and β10) and four α-helices (α1-α3, α6). The C-terminal domain (107-211) consists of four βstrands (β6-β9) along with two α-helices (α4 and α5) giving rise to a mixed αβ-sandwich arrangement and a long loop (L8, 156-179). Two short loops L4 (103-106) and L10 (212-215) connect the two domains and act as hinge regions for the domain movements. Mtb-DapB binds two ligands, a cofactor either NADH or NADPH and a cyclic substrate. The ADP part of NADH/NADPH is embedded in a solvent exposed groove like region located in the N-terminal domain and extending to the hinge region, while the nicotinamide part is placed in the floor of a relatively less exposed cavity C1 (Fig 3(B)) formed by residues from both N- and C-terminal domains. C-terminal side of C1 binds the substrate dihydrodipicolinate which is located near the nicotinamide ring of NADH/NADPH placed in the N-terminal side of C1. Nine experimental structures of Mtb-DapB have been reported in PDB. Most of these crystal structures and literature suggest both NADH and NADPH can bind in the cofactor binding site of the N-terminal region. A substrate mimetic competitive inhibitor 2,6-PDC (PDC) is bound at C1. In this study, we considered three crystal structures of Mtb-DapB, namely, 1P9L, 1C3V and 1YL7 as they are high resolution structures, the binding regions being occupied by NADH/NADPH and PDC so that all the ligand interaction features present in the binding sites can be thoroughly explored. Six different model systems were created from these PDB structures as narrated in the methodology section in order to represent various states of the protein in the cell, bound to one or more cofactors in presence and absence of the substrate molecule. With respect to the several stages in the reaction cycle, the geometric dynamic properties are quite likely to be different when the protein binds to different ligands and hence one needs to consider all these conformational states during ligand design. The hybrid molecules in (D-NDPHyb) and (D-NAIHyb) were designed in order to compare the stabilities of the interactions made individually by the ligands when they are free and when they are linked together with a linker. The stable interactions formed by the hybrid molecule can be mimicked to design inhibitors, which can compete for the binding sites of both the ligands of Mtb-DapB.

**Figure 3.**
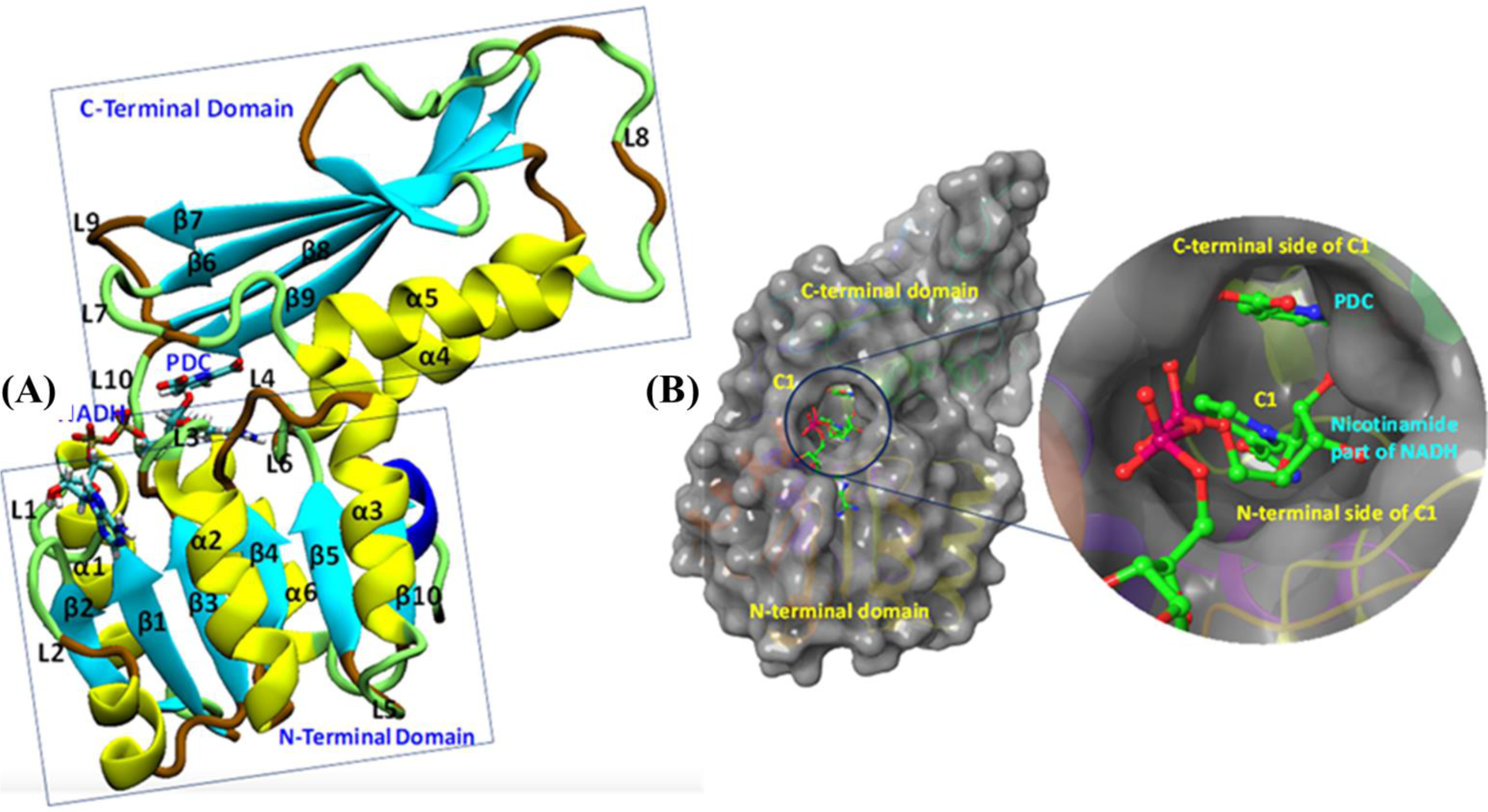
Mtb-DapB structure. **(A)** Overall structural architecture of Mtb-DapB, **(B)** NADH and substrate binding regions with a detailed picture of the C1 cavity.

### 3.2 Comparative analysis of the structural dynamics of Mtb-DapB model systems

The six model systems used in this study are based on similar initial structures, and provide a robust base to study how the binding of different ligands in different combinations affects the overall dynamics of Mtb-DapB and what conformation/s of which complex would be best to be used for drug design. Hence, various structural properties were analyzed from the MD trajectories of the six model systems using the ‘Simulation event analysis’ and ‘Simulation interaction diagram’ tools and we observed significant conformational variances in the binding sites of Mtb-DapB when bound to different ligands. Figure 4 shows the root mean squared deviations (RMSD) of the protein, root mean squared fluctuations (RMSF) of the Mtb-DapB residues RMSD, radius of gyration and solvent accessible surface areas (SASA) of the ligands in the six model systems.

**Figure 4.**
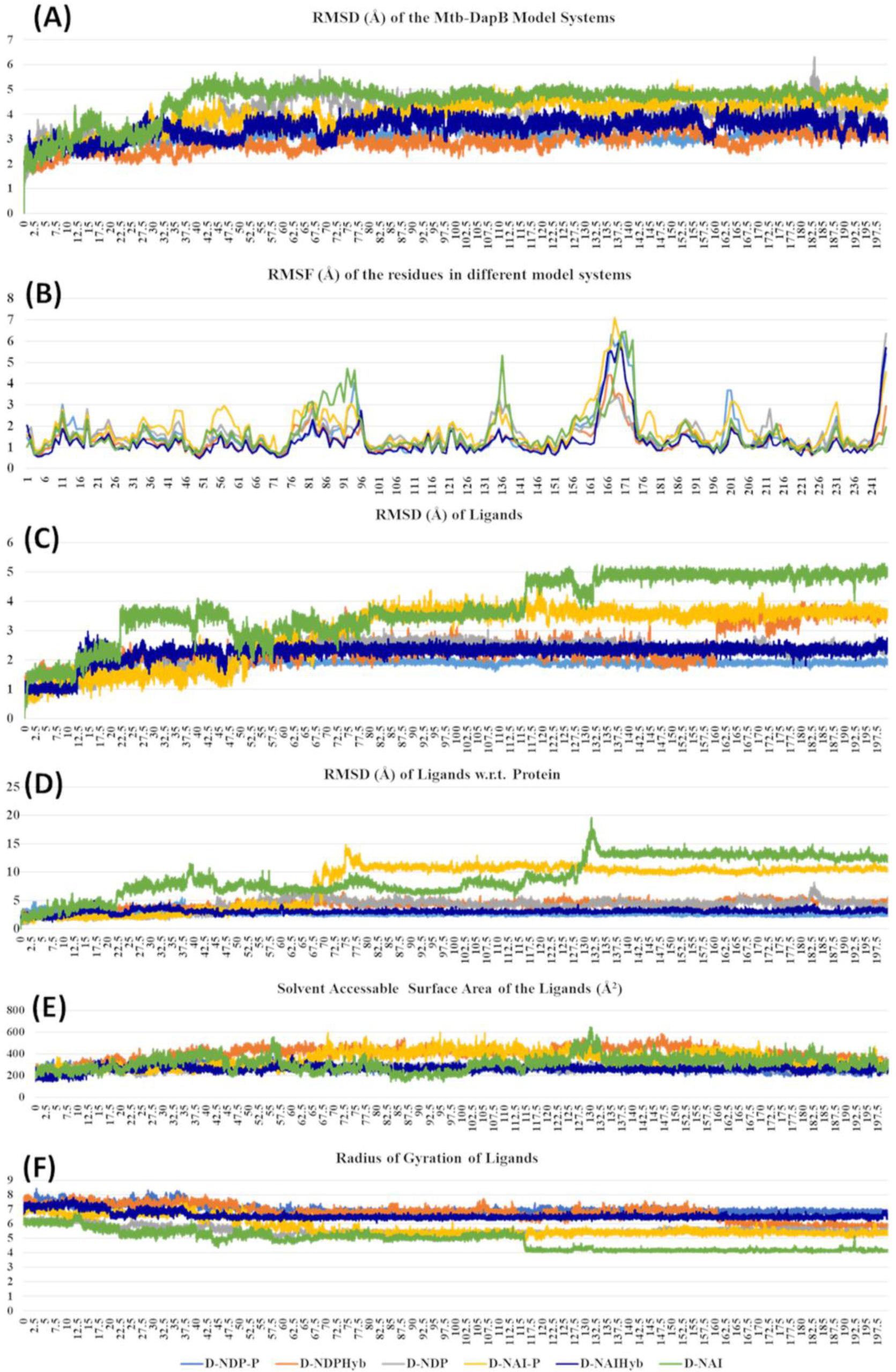
**(A)** RMSD of the protein, **(B)** RMSF of the Mtb-DapB residues, **(C)** RMSD of the ligands, **(D)** RMSD of the ligands w.r.t the protein, **(E)** Radius of gyration of the ligands **(F)** SASA of the ligands in the six model systems.

All the trajectories were well equilibrated by the end of 200 ns as observed from the RMSD graph. Though the initial structures of all the model systems are very similar, the model systems with only NADPH/NADH as in D-NDP and D-NAI show high RMSD from the respective initial structures as compared to the ones that additionally bind PDC in C1. The model systems where both the ligands were linked together to form hybrid molecules (D-NDPHyb and D-NAIHyb) showed lesser RMSD from their respective initial structures as compared to the ones where the ligands are separate (D-NDP-P and D-NAI-P) suggesting relatively stable complexes with the hybrid molecules. Also, the model systems binding NADH showed higher deviations from the initial structure except for D-NAIHyb. A quick look into the RMSD profiles of the ligand/s (NADPH/NADH and PDC considered together in case of D-NDP-P and D-NAI-P) with respect to themselves and with respect to the protein show high fluctuations in model systems D-NDP-P, D-NDP, D-NAI-P and D-NAI. These graphs suggest that NADH undergoes a drastic conformational change in D-NAI-P after 65ns and then remains stable till 200ns, while it almost detaches from the cavity after 130 ns in D-NAI. This observation is strengthened from radius of gyration and solvent accessible surface areas of the ligands throughout the simulations showing high fluctuations of NADH in D-NAI. As revealed from the C terminal L8 residues show higher (3.5 to 7.5 Å) fluctuations in all systems, which is however not a part of the binding pockets. In system D-NAI, the hinge region residue K136 shows high fluctuation.

### 3.3 Comparative analysis of stability of protein-ligand interactions in Mtb-DapB model systems

The simulation interaction diagram from Desmond gives the summary of all stable interactions between the active site residues and the ligands that last for more than 30% of the simulation. Figure 5 shows the fractions of different types of interactions made by the ligands with Mtb-DapB in the model systems and the stable interactions that lasted more than 30% of the simulation time for D-NAIHyb. Stable interactions for other model systems may be seen in Figure S1 and S2. The phosphate groups of the ADP part of NADPH in D-NDP-P make stable H-bonds with positively charged residues K9 and R14. The phosphate groups also make stable water bridges with F52 and T53 with occupancies of 80% and 60%, respectively. The –NH2 and -C=O groups of nicotinamide part make the most stable H-bonds with G75 (98%) and F105 (93%) as donor and acceptor, respectively. The –OH group of the sugar moiety attached to the nicotinamide ring makes stable H-bond with T77. Intramolecular H-bonds were also formed between the sugar –OH and the adjacent phosphate groups. In absence of PDC as in D-NDP, the Adenine base of NADPH makes H-bonds with H54 (58%) and N61 (36%). The –OH and the phosphate groups attached to the sugar moiety adjacent to the adenine base make water bridged interaction and H-bond with D33 (37%) and K9 (44%), respectively. The other phosphate group makes H-bond with F52 (42%). The sugar adjacent to the nicotinamide ring makes stable H-bonds with T33 (79%) and a water bridge interaction with T77 (64%). The H-bond interactions made by the –NH2 and -C=O groups of nicotinamide groups are same as those in D-NDP-P. The nicotinamide ring showed π-stacking with F17 as the latter is accessible due to absence of PDC. It was observed that many of the interactions formed by NADPH in D-NDP have occupancies <50% whereas the interactions formed in D-NDP-P showed better occupancy suggesting relatively stable binding of NADPH to Mtb-DapB in presence of PDC. NADH shows relatively unstable binding with Mtb-DapB as compared to NADPH both in presence and absence of PDC. The adenine part of NADH in D-NAI-P makes H-bond interactions with S211 (40%) and a cation-π interaction with R214. The phosphate groups make interactions with positively charged R214 and K11. Nicotinamide –NH2 and –C=O groups still make H-bond and water bridge with G75 and F105, respectively. However, except the interactions of R214, none of the other interactions showed more than 50% occupancy suggesting a weaker binding. In absence of PDC in D-NAI, NADH showed no stable interaction with Mtb-DapB as it showed tendency to move out of the binding pocket. PDC showed many stable interactions with the C1 residues in presence of both NADPH and NADH. In D-NDP-P, PDC makes H-bond/electrostatic interactions with the hinge region residue K136 (two, 98% and 93%), R214 (95%), T143 (two, 94% and 93%) G142 (45%), T77 (40%), H133 (54%), water bridges with P103 and D138 and a π-stacking with H132 (56%). Similarly, in D-NAI-P, PDC makes H-bond/electrostatic interactions with K136 (two, 88% and 83%), H133 (69%), water bridges with D138, H132 and N104 and makes a π-stacking with H132 (40%). The number and stability of interactions made by PDC was observed to be higher in presence of NADPH than NADH. From these observations of interactions made by NADPH/NADH and PDC, we also get a hint that the strength of cofactor binding is higher in presence of the substrate. Another important observation is the interactions made by the hybrid molecules, phosphate groups of the hybrid molecule formed by linking NADPH and PDC (NDPHyb) made stable H-bond/electrostatic interactions with K509 (45%), R214 (two, 65% and 53%), the nicotinamide part made π-stacking with F52, and H-Bonds with F105 (45%) and A102 (60%). The PDC part of NDPHyb retained its stable interactions in the D-NDPHyb model system with K136 (99%), H133 (48%), T143 (two, 86% each), S141 (41%), G142 (75%) and water bridges with D138 (93%). It was interesting to observe that, when NADH was linked with PDC to form a hybrid molecule (NAIHyb), it formed more number of stable interactions with Mtb-DapB owing to its structural stability, which was not formed in D-NAI-P or D-NAI.

**Figure 5.**
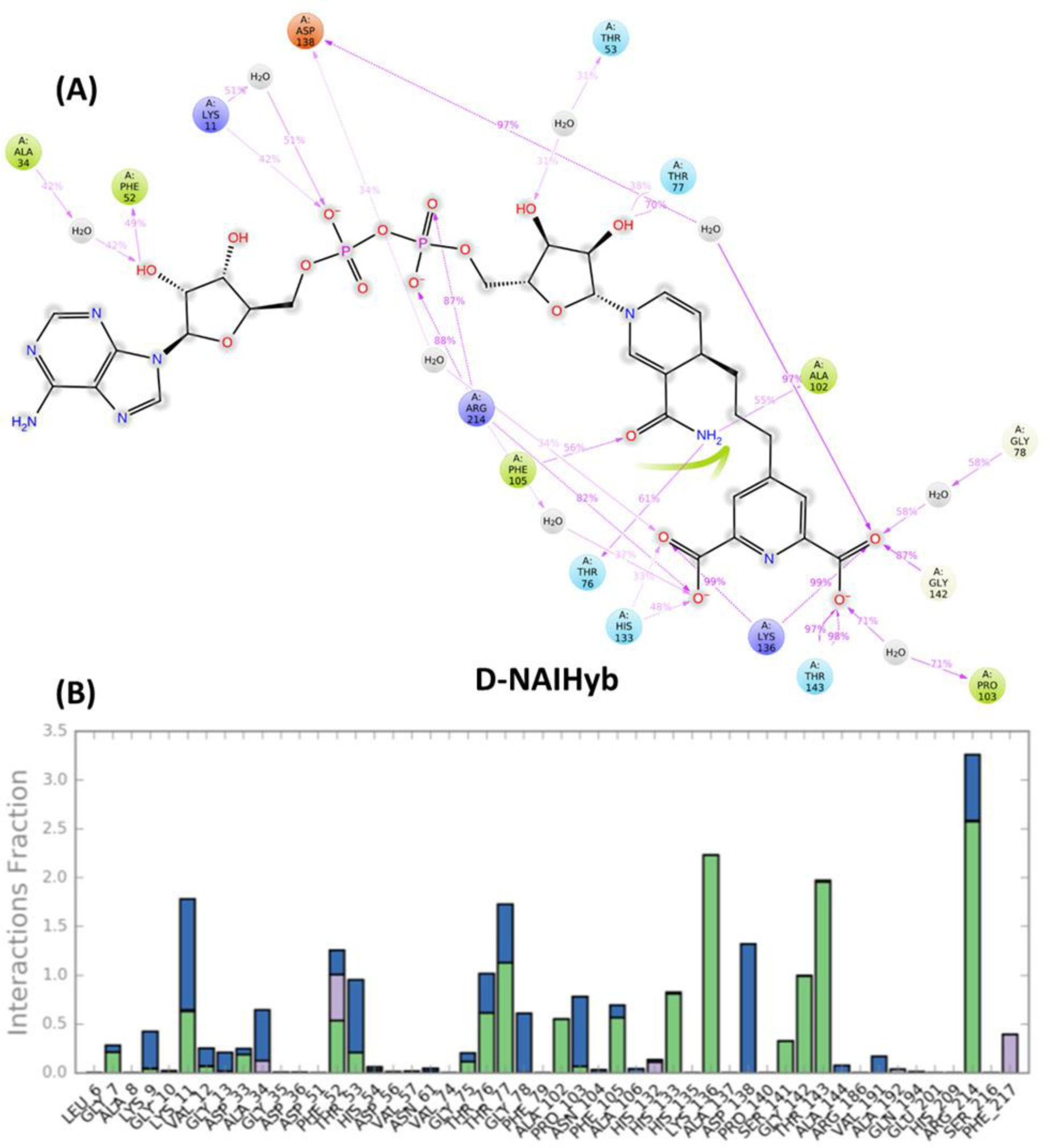
**(A)** The stable interactions that lasted more than 30% of the simulation time, **(B)** Fractions of different types of interactions made by the ligands with *Mtb*-DapB in the model system D-NAIHyb as an example.

The sugar moiety attached to the adenine part of NAIHyb makes H-bond interactions with F52 (49%) and water bridge with A34 (42%). The phosphate groups make H-bond/electrostatic interactions with K11 (42%) and R214 (two, 88% and 87%) while the sugar attached to the nicotinamide ring makes H-bond with T77 (70%). The n-NH2 and –OH groups of nicotinamide ring H-bond with T76 (61%), A102 (55%) and F105 (56%) respectively. The PDC part of NAIHyb retains its strong H-bond/electrostatic interactions with K136 (two, 99% each), T143 (two, 98%, 97%), G142 (87%), H133 (48%) and water bridges with D138 (97%), P103 (71%) and G78 (58%).

Thus, the hybrid molecules form stable protein-ligand complexes by forming additional interactions with the Mtb-DapB binding site, especially in the C1 region. From the above observations it is clear that in all model systems, the interactions formed by the C1 residues were relatively more stable than the ones formed by other parts of the binding site. This observation was considered while generating hybrid pharmacophore models, which is discussed in the next section.

### 3.4 Dynamics based conventional and hybrid pharmacophore models

To explore interaction features of binding sites in different conformational states, each of the six MD trajectories were clustered using hierarchical clustering based on mutual RMSD to obtain representative conformations which have RMSD of least 3Å with respect to each other. From D-NDP-P, D-NDP, D-NDPHyb, D-NAI-P and D-NAIHyb model systems 3, 7, 3, 3 and 2 clusters were obtained respectively. DapB-NAI system was not considered for pharmacophore modelling as none of the interactions made by NADH in this model system showed >40% occupancy. The cluster representatives were further verified if they are showing all the stable interactions as identified from the MD simulations. We observed that 2, 3, 2, 1 and 2 cluster representatives from the MD trajectories of D-NDP-P, D-NDP, D-NDPHyb, D-NAI-P and D-NAIHyb respectively were able to show all stable interactions and were used to generate the N- and H-type pharmacophore models. Apart from the 4 representatives from D-NDPHyb and D-NAIHyb, the conventional models generated from the 2 representatives of D-NDP-P were combined to generate 2 H-type models. Details of six H-type and six N-type models are described in the Supplementary tables S1 and S2.

Figure 6 shows one representative from H- and N-type pharmacophore models and Tables S1, S2 and Figures S3 and S4 give the details of all the models belonging to the three categories. All the models consisted of 5 to 8 features. The H-type models were designed to represent the stable interactions made by both PDC and the cofactor, excluding the ADP region. The reasons for excluding the ADP region are as follows, 1) ADP is a common metabolite in both human and Mtb, so mimicking its interaction features might cause specificity issue 2) combination of strong interaction features from both the cofactor and substrate might fetch chemically diverse molecules with good binding affinity and 3) interactions of the cofactors, substrate and PDC were relatively stable with the C1 residues (ADP region of the cofactors/hybrid molecules do not bind near C1). H-type pharmacophore models were obtained by two different ways. First, from the interactions of the Hybrid constructs (excluding the ADP part) as in D-NDPHyb and D-NAIHyb and second way was by merging the conventional models from NADPH/NADH and PDC of D-NDP-P followed by eliminating the features obtained from the ADP part of NADPH/NADH (Figure 1).

**Figure 6.**
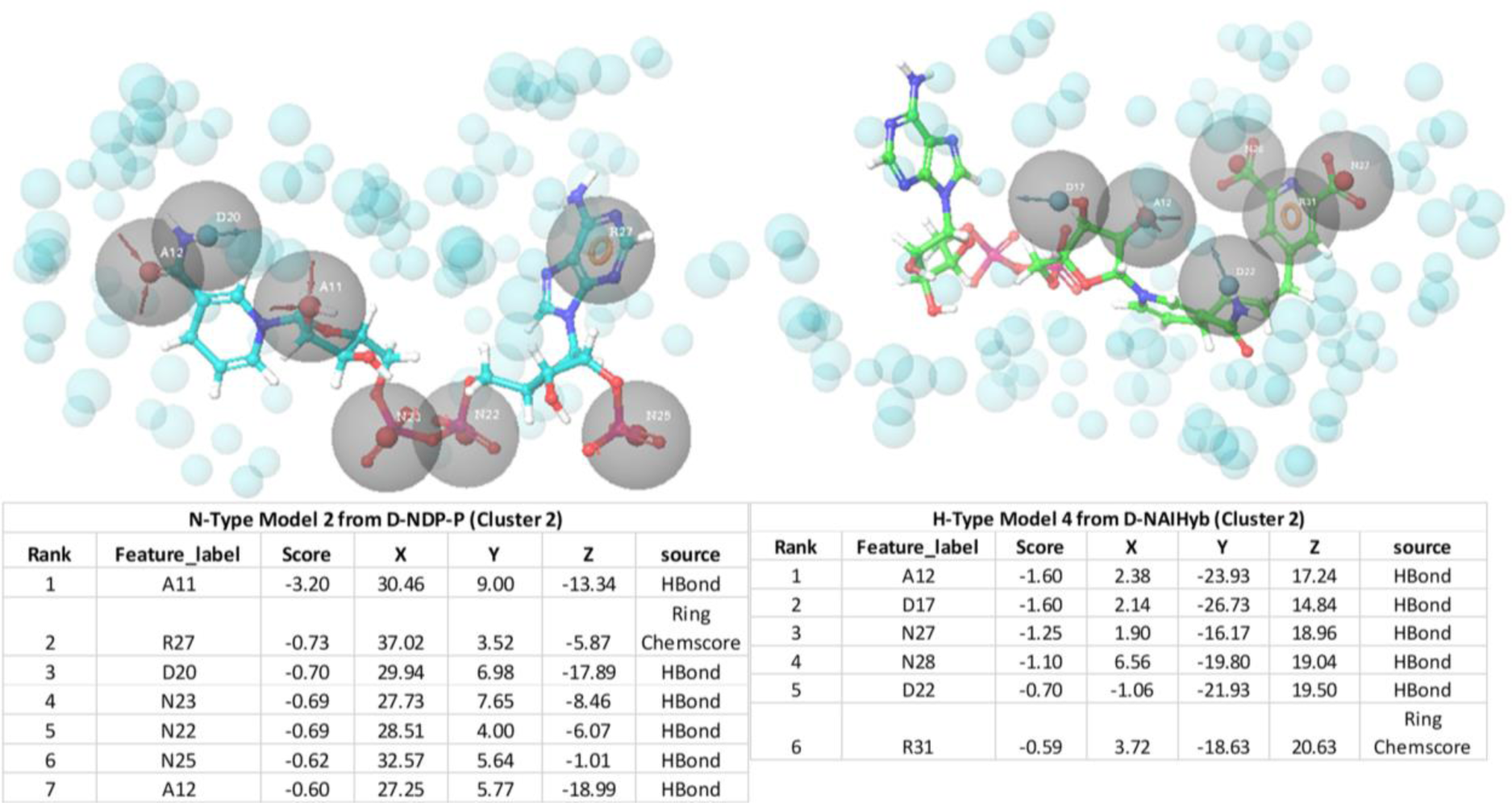
Snapshots of one example from N-type and H-type pharmacophore models, with their respective feature tables.

To validate the performances of our pharmacophore models, known Mtb inhibitors are obtained from ChEMBL. As mentioned before, only a few inhibitors of Mtb-DapB have been reported in literature and these product analogue inhibitors (Singh et al., 2013) and sulphonamide inhibitors (Paiva et al., 2001) show IC50 > 26 μM and Ki values ranging from 7 to 48 µM respectively. Secondly, our pharmacophore models are based on the interactions of the cofactors (N-type) and the cofactor-substrate hybrid molecules (H-type) with Mtb-DapB. No DapB inhibitor acting through interaction with both of the binding site residues is reported yet. Hence, we selected compounds from ChEMBL with low MIC values (≤ 1μM) as the test set to evaluate the model performance. A total of 4298 molecules with MIC ≤ 1μM reported in ChEMBL database and 155 Mtb active molecules reported by GSK were subjected to structure similarity-based clustering. The molecules clustered with Tanimoto similarity **≥** 0.85 were eliminated and the rest 2349 molecules were used as the active dataset. In addition, 2934 Mtb inactive compounds are also selected from ChEMBL (reported as ‘inactive’ or with MIC ≥ 50μM). A total of 12934 molecules including 10000 random decoy molecules are obtained from DUD.E database and along with 2934 Mtb inactive molecules from ChEMBL were taken as the inactive data set. Hypothesis validation program of Phase module of Schrodinger suite was used to screen both the active and inactive datasets against all the 6 H-type and 6 N-type models. Interestingly, it was observed from the percentage screen plots that, the H-type models were able to screen more number of Mtb-active compounds as compared to the N-Type models. Though the percentage of Mtb hits by the H-type models are very low (10-15%), but they are ranked high in the list of hits. And considering that these are not exactly active on Mtb-DapB (as target is not reported), but they are active on Mtb whole cells, it can be safely concluded that the models represent Mtb specific interaction features, which are not available in a reasonably large random decoy library. The ROC curves (Figure S5) for these hypothesis validation screening, are not near the random line. The numbers of active and inactive compounds screened by each H and N-type model is given in Table S3. For these sets of Mtb-active and decoy molecules, the H-type pharmacophores based on the interaction features of NADPH-PDC showed better performance to screen Mtb-active molecules as compared to the H-type models based on NADH-PDC.

The ligand dataset curated from publicly available chemical databases was then screened by these N- and H-type models and molecules that matched at least 5 features were retained as hits. The unique consolidated list of compound hits screened by the N- and H-type models were named as N-set and H-set respectively. N-Set contained 2884 molecules while the H-set contained 3707 molecules. Figure S6 and S7 show the pharmacophore models mapped to the respective top hits with best fitness scores.

### 3.5 Comparative analysis of N-set and H-set compounds

As the aim of this study is to explore the hybrid pharmacophore model as an effective tool to screen new chemotypes with better affinities with DapB as compared to the conventional models, we compared different aspects of N-set and H-set compounds, such as their structural similarities, their binding energies with Mtb-DapB and drug like and ADMET properties.

#### 3.5.1 Comparison of Mtb-DapB binding affinities

In order to quantify the binding affinities of the H-set and N-set compounds with Mtb-DapB, all the 3707 H-set and 2884 N-set compounds were docked with each of the 10 representative structures (described in the ‘Dynamics based conventional and hybrid pharmacophore models’ section) used to generate the pharmacophore models. The best scoring pose for each compound among the 10 docking calculations was retained. This was followed by MMGBSA binding free energy calculations taking all these protein ligand complexes. Figure 7(A) shows the distributions of the XP docking scores and the MMGBSA ΔGbind ligand efficiency values of the top 1000 H-set and the top 1000 N-set compounds. The graphs show that, 72.3% of the top 1000 H-set compounds had docking scores below -7, while only 34% of the N-set compounds have this range of XP-docking score. 20.1% of the top scoring 1000 H-set compounds had docking score <-8, while only 7.7% of the N-set compounds had this score. The MMGBSA ΔGbind ligand efficiency values were obtained by normalizing the MMGBSA ΔGbind by the number of heavy atoms present in the respective compounds. Figure 7(A) shows that 80.9% of the top 1000 H-set compounds showed ligand efficiency above 2.2, while only 40.4% of the top 1000 N-set compounds showed this range. These comparative graphs clearly indicate that the H-set compounds have better binding affinities with Mtb-DapB as compared to the N-set compounds. Examples of the best scoring H-set and N-set compounds have been shown in Figure 7(B).

**Figure 7.**
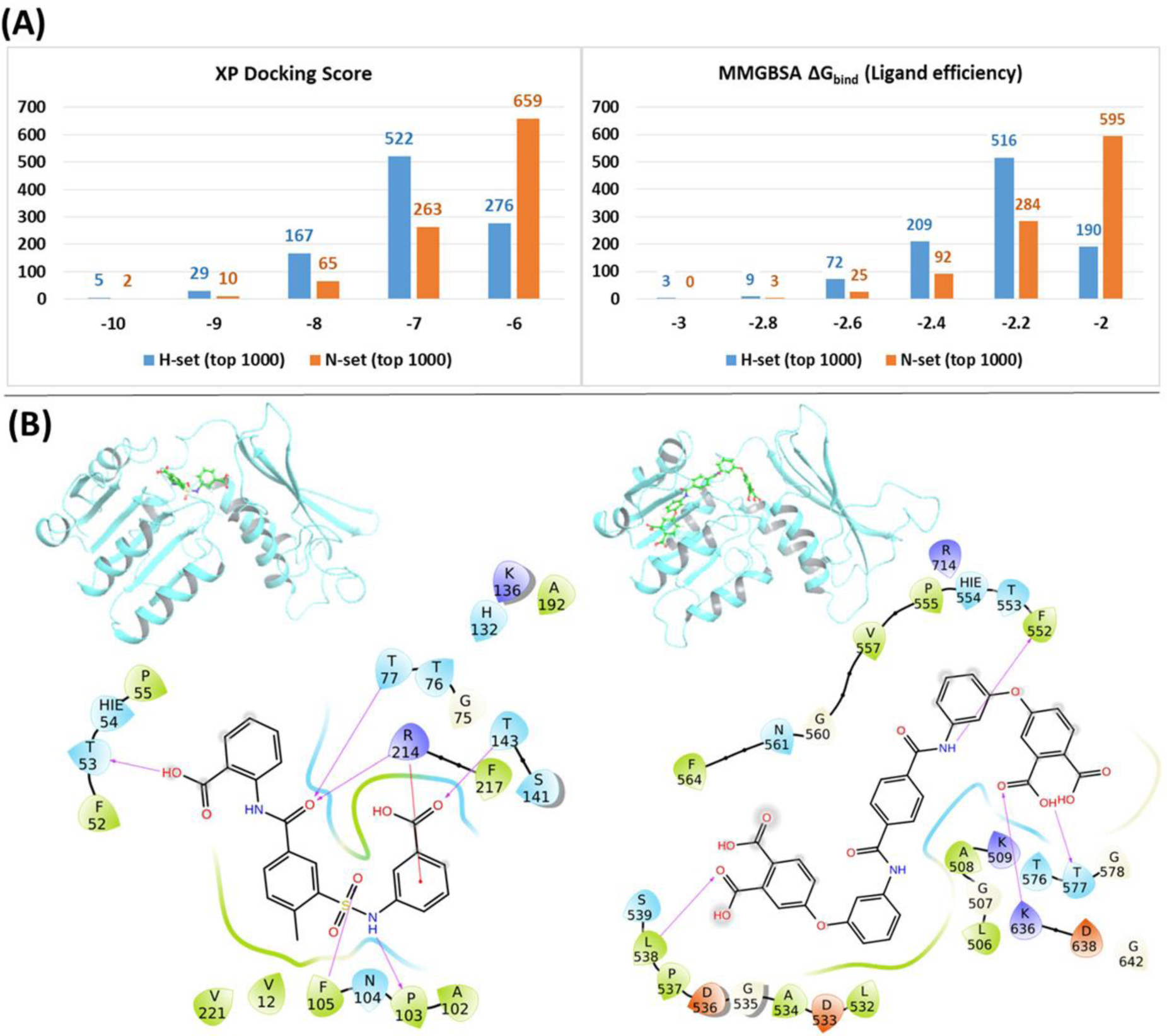
**(A)** Distributions of the XP docking scores and the MMGBSA ΔGbind ligand efficiency values of the top 1000 H-set and the top 1000 N-set compounds, **(B)** Examples of interactions of the best scoring H-set and N-set compounds bound to the respective binding sites.

#### 3.5.1 Comparison of structural features

The structures of the H-set compounds were compared with those of the N-set compounds using different measures. The H-set compounds were chosen as the query library and compared against N-set as the reference library. For each compound in the query library, the nearest neighbor in the reference library was obtained using fingerprint similarities based on different matrices. Four different types of binary hashed fingerprints such as linear, radial, MOLPRINT2D and atom triplets were calculated for the two libraries. The nearest neighbor similarities were obtained based on different similarity matrices, viz., Tanimoto, Cosine, Dice and Tversky similarity matrices and plotted as histograms. Figure S8 shows the similarity score distributions between H-set vs. N-set. Histograms in Figure 8(A) clearly indicate that more than 70% of the H-set compounds have scores below 0.6 with respect to the N-set compounds. This shows that the hybrid pharmacophore models screen structurally different molecules as compared to the conventional models.

**Figure 8.**
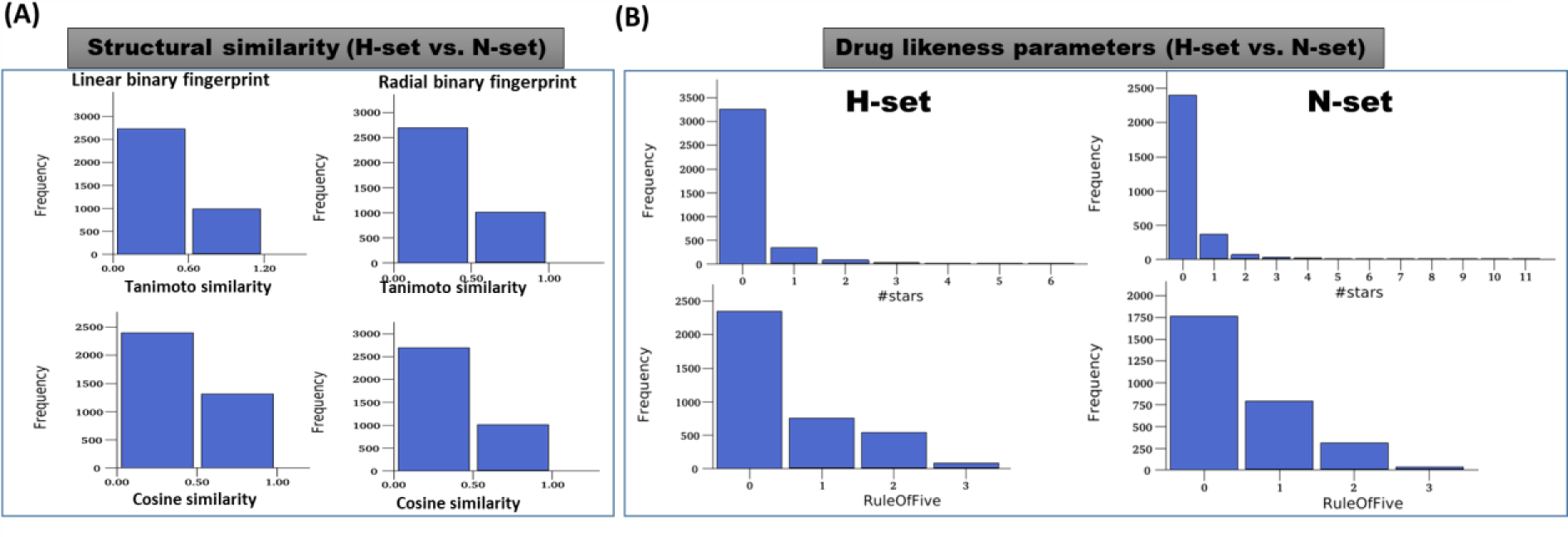
**(A)** Distribution of the maximum similarity (MaxSim) scores of the H-set compounds with respect to N-set compounds based on the linear and radial fingerprints. MaxSim represents the similarity score (calculated using Tanimoto and Cosine matrices) of each molecule from H-set with the nearest neighbor of the N-set. Distribution of this MaxSim scores for the molecules of one set with respect to the other set gives an idea about the structural similarities between the libraries (B) Histograms of important drug likeness parameters of the H-and N-set compounds.

#### 3.5.3 Comparison of druglike properties

Various drug likeness parameters of the H-set and N-set compounds were calculated using QuikProp and the drug like properties of both sets were compared to each other. Figure 8(B) shows the histograms of various drug likeness parameters of the H- and N-set compounds. The solubility, bioavailability and the drug likeness scores were found to follow strikingly different trends in case of N and H set compounds. Figure S9 shows distribution of different druglike properties of the H-set and N-set compounds. The #star descriptor indicates the number of property or descriptor values (such as molecular weight, dipole moment, ionization potential, electron affinity, SASA and its components, volume, HB donor and acceptor, globularity, solubility, lipophilicity, bioavailability, toxicity etc. (refer List S1 for detailed information)) that fall outside the 95% range of similar values for known drugs. A higher value range (about 40% molecules have a value >=5) of #stars for the N-set compounds suggests they are less drug-like than H-set molecules with few stars (only 1% molecules have a value >=5). Similarly, the #RO5 values (Number of violations of Lipinski’s rule of five) are also higher for the N-set compounds as compared to the H-set compounds (Figure 8 (B)) showing better drug likeliness of the later. The #metab descriptor is a predicted value representing number of likely metabolic reactions gives an estimation of the off-target interactions and toxicity of the compounds. The N-set compounds show a higher number as compared to the H-set compounds. Hence with all these comparative observations of the structural and physicochemical properties, and drug-likeliness scores, we can summarize that, the hybrid pharmacophore models lead to a structurally diverse and more druglike chemical space as compared to the conventional models.

## 4 Conclusions

The present study reports a robust computational approach, wherein, six different model systems of Mtb-DapB binding different combinations of ligands were modelled and each of them were subjected to 200 ns molecular dynamics simulations. The structural and enthalpic stabilities of these model systems were monitored throughout the simulations and it was revealed that the hybrid ligands designed by linking two native ligands (cofactor and substrate) of Mtb-DapB are able to make highly stable non-covalent interactions in the binding pockets. These stable interactions formed by the hybrid ligands with two adjacent binding site regions of Mtb-DapB were utilized to generate hybrid dynamics-based pharmacophore models. The abilities of these hybrid models to screen molecules with new chemotypes, better binding affinities, and drug-like properties were comparatively assessed with that of the conventional models generated from the native ligands. Cheminformatics based structure comparison, docking scores, binding energies and ADMET properties of the molecules screened by the hybrid pharmacophore models were found to be more druglike, thus establishing the hybrid models as efficient tools to venture into novel anti-TB chemical space.

## 5 Supporting Information

Supporting information (file type: PDF) containing the following data is available free of charge.

- Section S1. Details of MD simulation methodology
- Figure S1. Stable protein-ligand interactions in the six model systems that lasted more than 30% of the simulation time.
- Figure S2. Fractions of different types of interactions made by the ligands with Mtb-DapB in the six model systems.
- Table S1. Feature tables of all the six N-type pharmacophore models.
- Table S2. Feature tables of all the six H-type pharmacophore models.
- Figure S3. Snapshots of N-type pharmacophores with the cofactors aligned to them.
- Figure S4. Snapshots of H-type pharmacophores with the first four aligned to the respective hybrid molecules. The last two are designed by combining two pharmacophore models, hence do not have a reference molecule.
- Table S3 Screening results of Mtb-active and decoy molecules
- Figure S5. ROC curves for Mtb-active and decoy molecules screened again H-type pharmacophore modelsFigure S6. N-type pharmacophore models mapped to the respective top scoring compounds.
- Figure S7. H-type pharmacophore models mapped to the respective top scoring compounds.
- Figure S7. The similarity score distributions between H-set vs N-set
- Figure S9. Distribution of different pharmacokinetic properties of the H-set and N-set compounds.
- List S1. Details of all types QuikProp descriptors calculated in this study (recommended range of value for each property is mentioned in brackets).

## 6 Author Contributions

AB and CC conceived the project. CC performed calculations and AB and CC analyzed the results. AB and CC prepared the manuscript. Both authors read and approved the final manuscript.

## 9 Acknowledgements

AB and CC thank Schrodinger Inc. for providing short term license for some modules and Prof. Prasad V. Bharatham of National Institute of Pharmaceutical Education and Research, Mohali, for allowing to use software resources at his lab.

