## Supplementary File for "Hybrid dynamic pharmacophore models as effective tools to identify novel chemotypes for anti-TB inhibitor design: A case study with Mtb-DapB"

Chinmayee Choudhury[1],

Anshu Bhardwaj[2*]

[1]Department of Experimental Medicine and Biotechnology, Post Graduate institute of Medical Education and Research, Sector 12, Chandigarh, India

[2]Bioinformatics Centre, CSIR-Institute of Microbial Technology, Sector 39A, Chandigarh, India

*Corresponding author

**Section S1.** Details of MD simulation methodology

**Molecular Dynamics (MD) Simulations**

The Protein Preparation Wizard (PPW) module (Shivakumar et al., 2010) of Schrödinger software package, version 2019-2 was used to pre-process the macromolecular structure downloaded from PDB. Missing hydrogens were added, water molecules beyond 5 Å of the active site were removed and appropriate bond orders were assigned to the structure. The residues/side chains unresolved in some of the crystal structures were repaired with prime (Jacobson et al., 2002) module in the PPW. The protonation states of the polar residues were optimized with the protassign module of PPW, which uses PROPKA to predict pKa values (pH 7.0±2.0) and side chain functional group orientations. The structures were then subjected to restrained minimization (cutoff RMSD 0.3 Å) with impref to avoid steric clashes. These structures were further used for molecular dynamics (MD) simulations. MD simulations were carried out on the six model systems using the Desmond MD simulation package (release 2017) (Shaw et al., 2014). The OPLS_2005 (Jorgensen and Tirado-Rives, 1988) force field was employed for the protein-ligand complexes. Using the system builder tool of DESMOND the six model systems of Mtb-DapB were solvated in a cubical water box (TIP3P water model) keeping 10 Å buffer space in x, y and z dimensions. Each system was neutralized by adding appropriate counter ions and an ionic concentration of 0.15 M was maintained by adding Na+ and Cl− ions. The systems were minimized with 10000 steepest descent steps followed by gradual heating from 0 to 300 K, under NVT ensemble. The systems were thermally relaxed before the production run using Nose-Hoover Chain thermostat method for 1 picosecond and 1 picosecond of pressure relaxation with Martyna-Tobias-Klein barostat method. Finally, 0.2 microsecond (200 ns) production run under NPT ensemble was carried out for each system using a cutoff distance of 12 Å for non-bonded interactions. Simulation quality analysis, simulation event analysis, simulation interaction diagrams were used for trajectory analysis.

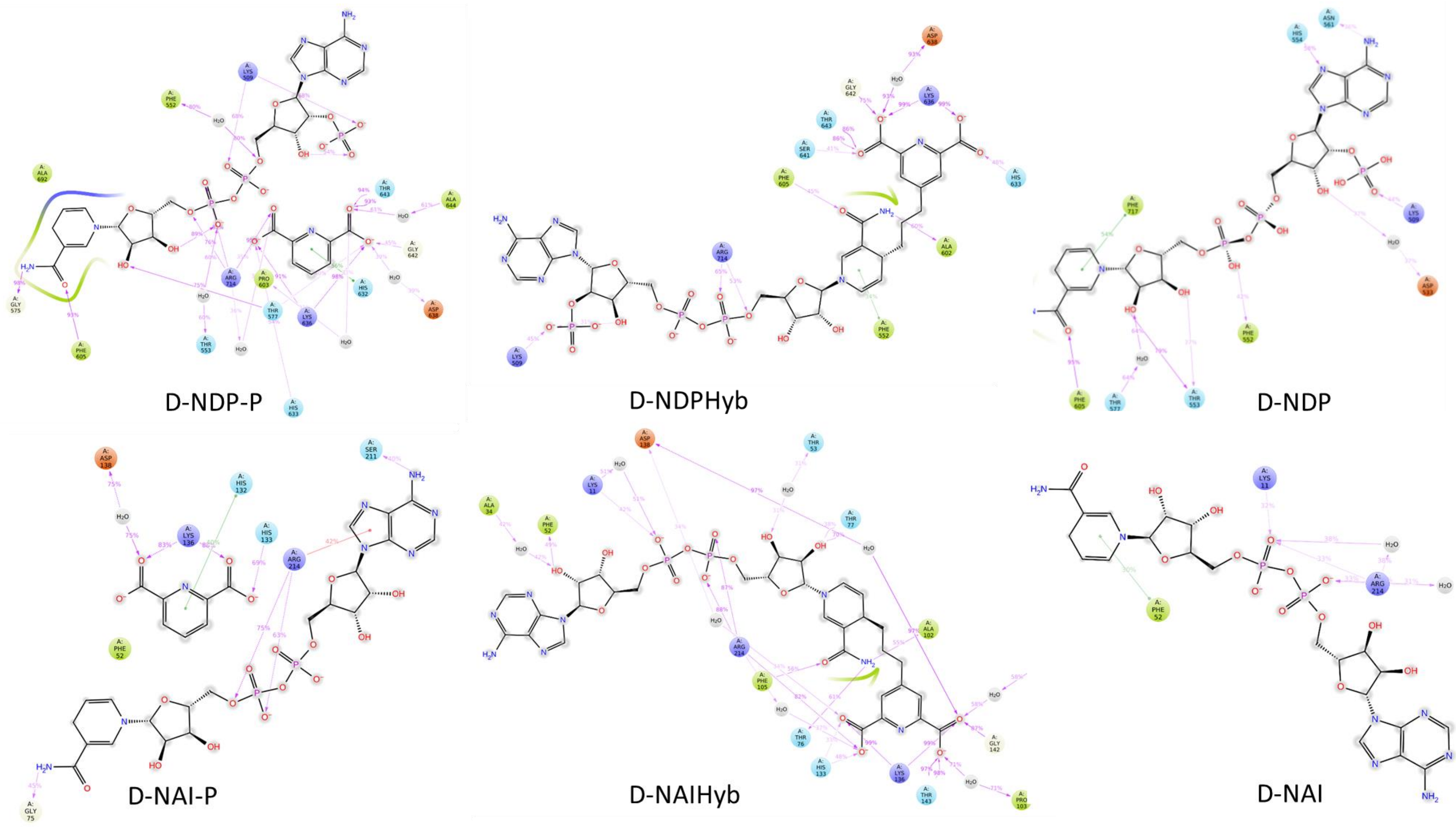


**Figure S1.** Stable protein-ligand interactions in the six model systems that lasted more than 30% of the simulation time.

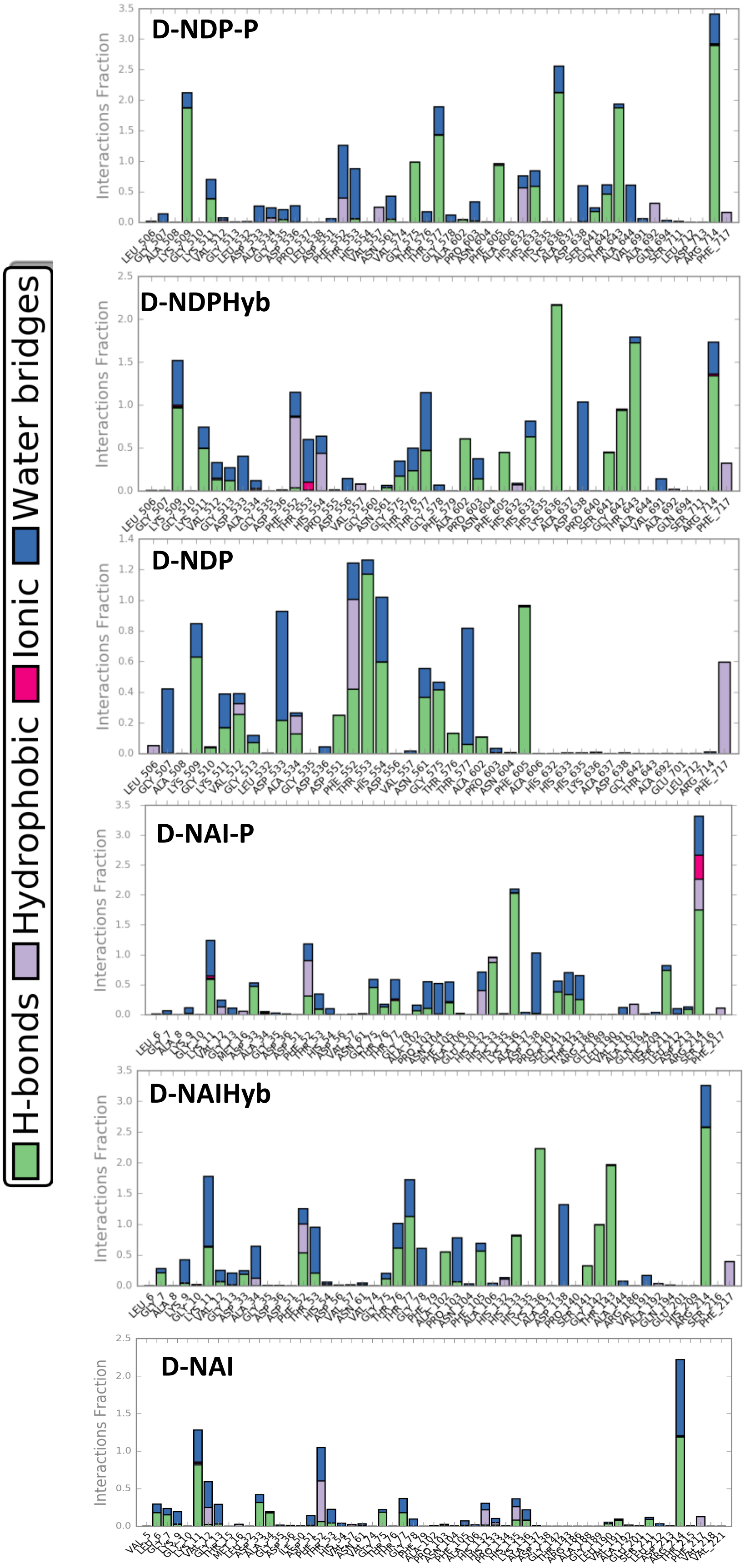


**Figure S2.** Fractions of different types of interactions made by the ligands with *Mtb*-DapB in the six model systems.

**Table S1.** Feature tables of all the six N-type pharmacophore models.

| | | | | | | |
|---|---|---|---|---|---|---|
| | | **N-Type Model1 D-NDP-P (Cluster 1)** | | | | |
| 1 | A11 | -1.6 | 13.7992 | 33.9899 | 18.0959 | HBond |
| 2 | N23 | -0.69 | 18.3601 | 32.1812 | 14.6208 | HBond |
| 3 | N22 | -0.69 | 20.3436 | 28.4824 | 14.9029 | HBond |
| 4 | R27 | -0.67 | 13.8428 | 21.9542 | 16.029 | RingChemscoreHphobe |
| 5 | D20 | -0.65 | 14.6187 | 34.3047 | 23.6665 | HBond |
| 6 | N25 | -0.62 | 18.3972 | 22.7516 | 12.3691 | HBond |
| 7 | A8 | -0.33 | 18.2548 | 31.3989 | 16.7327 | HBond |
| 8 | A12 | -0.23 | 17.2656 | 35.5536 | 24.7898 | HBond |
| | | **N-Type Model2 D-NDP-P (Cluster 2)** | | | | |
| 1 | A11 | -3.20 | 30.46 | 9.00 | -13.34 | HBond |
| 2 | R27 | -0.73 | 37.02 | 3.52 | -5.87 | Ring Chemscore |
| 3 | D20 | -0.70 | 29.94 | 6.98 | -17.89 | HBond |
| 4 | N23 | -0.69 | 27.73 | 7.65 | -8.46 | HBond |
| 5 | N22 | -0.69 | 28.51 | 4.00 | -6.07 | HBond |
| 6 | N25 | -0.62 | 32.57 | 5.64 | -1.01 | HBond |
| 7 | A12 | -0.60 | 27.25 | 5.77 | -18.99 | HBond |
| | | **N-Type Model3 D-NDP (Cluster 2)** | | | | |
| 1 | D23 | -1.6 | 25.1455 | 39.5989 | 40.6352 | HBond |
| 2 | R30 | -0.76 | 27.4781 | 34.7701 | 47.6988 | RingChemscoreHphobe |
| 3 | A12 | -0.7 | 19.4434 | 36.2985 | 36.3583 | HBond |
| 4 | D24 | -0.67 | 30.1244 | 33.9834 | 45.8677 | HBond |
| 5 | A1 | -0.08 | 27.9976 | 37.4251 | 46.8745 | HBond |
| | | **N-Type Model4 D-NDP (Cluster 4)** | | | | |
| 1 | D22 | -1.6 | 12.653 | 8.55 | 22.4881 | HBond |
| 2 | A19 | -1.33 | 12.6881 | 0.7402 | 29.684 | HBond |
| 3 | R30 | -0.85 | 17.7298 | 3.943 | 26.6276 | RingChemscoreHphobe |
| 4 | A12 | -0.7 | 8.0939 | 3.5405 | 15.7099 | HBond |
| 5 | D27 | -0.7 | 11.0348 | 4.5953 | 15.822 | HBond |
| | | **N-Type Model5 D-NDP (Cluster 6)** | | | | |
| 1 | A20 | -1.6 | 25.0936 | -7.0556 | -17.9801 | HBond |
| 2 | A14 | -1.03 | 19.5745 | -7.1139 | -10.2979 | HBond |
| 3 | D22 | -0.84 | 18.831 | -1.8391 | -12.4926 | HBond |
| 4 | R30 | -0.8 | 21.9953 | -5.7205 | -19.8722 | RingChemscoreHphobe |
| 5 | D25 | -0.7 | 20.6307 | -3.7192 | -21.9417 | HBond |
| 6 | A12 | -0.47 | 10.1189 | -4.0745 | -8.0887 | HBond |
| 7 | D27 | -0.16 | 11.0907 | -2.2234 | -10.3843 | HBond |
| 8 | D21 | -0.05 | 24.4512 | -8.312 | -14.9217 | HBond |
| | | **N-Type Model6 D-NAI-P (Cluster 1** | | | | |
| 1 | N24 | -1.91 | 5.0175 | 4.2839 | -6.2929 | HBond |
| 2 | D19 | -0.67 | 12.6955 | -0.3395 | -9.8709 | HBond |
| 3 | A2 | -0.58 | 12.4315 | 2.112 | -10.3629 | HBond |
| 4 | H22 | -0.52 | 9.883 | 10.3092 | -6.9477 | PhobEn |
| 5 | A8 | -0.29 | 5.9034 | 5.5617 | -7.9497 | HBond |
| 6 | R25 | -0.19 | 9.0993 | 2.2892 | -10.3039 | RingChemscoreHphobe |

**Table S2.** Feature tables of all the six H-type pharmacophore models.

| Rank | Feature_label | Score | X | Y | Z | source |
|---|---|---|---|---|---|---|
| **H-Type Model1 D-NDPHyb (Cluster 2)** | | | | | | |
| 1 | N28 | -1.85 | 18.82 | 46.6723 | -6.4479 | HBond |
| 2 | N29 | -0.75 | 23.3865 | 42.902 | -6.7673 | HBond |
| 3 | D20 | -0.7 | 13.9691 | 41.0351 | -1.9275 | HBond |
| 4 | N25 | -0.66 | 23.4173 | 38.1076 | -6.9987 | HBond |
| 5 | R32 | -0.46 | 20.6684 | 44.1271 | -4.9795 | RingChemscoreHphobe |
| 6 | A9 | -0.21 | 21.5766 | 38.6976 | -5.8601 | HBond |
| **H-Type Model 2 from D-NDPHyb (Cluster 3)** | | | | | | |
| 1 | A12 | -2.44 | -27.39 | -12.27 | -15.83 | HBond |
| 2 | N29 | -0.87 | -29.56 | -7.39 | -20.94 | HBond |
| 3 | N28 | -0.65 | -31.46 | -5.61 | -15.48 | HBond |
| 4 | D21 | -0.62 | -24.75 | -9.86 | -12.77 | HBond |
| 5 | A13 | -0.51 | -23.05 | -7.36 | -13.19 | HBond |
| 6 | R32 | -0.33 | -28.82 | -6.26 | -17.72 | Ring Chemscore |
| **H-Type Model 3 from D-NAIHyb (Cluster 1)** | | | | | | |
| 1 | N27 | -2.04 | 22.9181 | 24.1707 | 3.8691 | HBond |
| 2 | A12 | -1.6 | 24.0897 | 17.0952 | 8.524 | HBond |
| 3 | N28 | -1.2 | 19.9565 | 22.6982 | 8.763 | HBond |
| 4 | D22 | -0.62 | 23.4871 | 17.261 | 3.2131 | HBond |
| 5 | R31 | -0.55 | 20.219 | 22.5253 | 5.3252 | RingChemscoreHphobe |
| **H-Type Model 4 from D-NAIHyb (Cluster 2)** | | | | | | |
| 1 | A12 | -1.60 | 2.38 | -23.93 | 17.24 | HBond |
| 2 | D17 | -1.60 | 2.14 | -26.73 | 14.84 | HBond |
| 3 | N27 | -1.25 | 1.90 | -16.17 | 18.96 | HBond |
| 4 | N28 | -1.10 | 6.56 | -19.80 | 19.04 | HBond |
| 5 | D22 | -0.70 | -1.06 | -21.93 | 19.50 | HBond |
| 6 | R31 | -0.59 | 3.72 | -18.63 | 20.63 | Ring Chemscore |
| **H-Type Model5 D-NDP-P (Cluster 1)** | | | | | | |
| 1 | N5 | -1.87 | 13.7992 | 33.9899 | 18.0959 | HBond |
| 2 | A1 | -1.6 | 14.6187 | 34.3047 | 23.6665 | HBond |
| 3 | N6 | -1.6 | 18.2548 | 31.3989 | 16.7327 | HBond |
| 4 | D4 | -0.65 | 17.2656 | 35.5536 | 24.7898 | HBond |
| 5 | R7 | -0.37 | 26.0106 | 13.0689 | -17.513 | RingChemscoreHphobe |
| 6 | A2 | -0.33 | 23.2091 | 10.3234 | -13.0278 | HBond |
| 7 | A3 | -0.23 | 24.4345 | 10.21 | -16.2898 | HBond |
| **H-Type Model6 D-NDP-P (Cluster 2)** | | | | | | |
| 1 | A1 | -3.2 | 30.46 | 8.9999 | -13.339 | HBond |
| 2 | N4 | -2.18 | 20.963 | 36.9744 | 16.5854 | HBond |
| 3 | N5 | -1.36 | 15.4363 | 39.1187 | 17.9083 | HBond |
| 4 | D3 | -0.7 | 29.9351 | 6.98 | -17.8893 | HBond |
| 5 | R6 | -0.66 | 18.9053 | 38.8579 | 18.71 | RingChemscoreHphobe |
| 6 | A2 | -0.6 | 27.2546 | 5.7702 | -18.9887 | HBond |

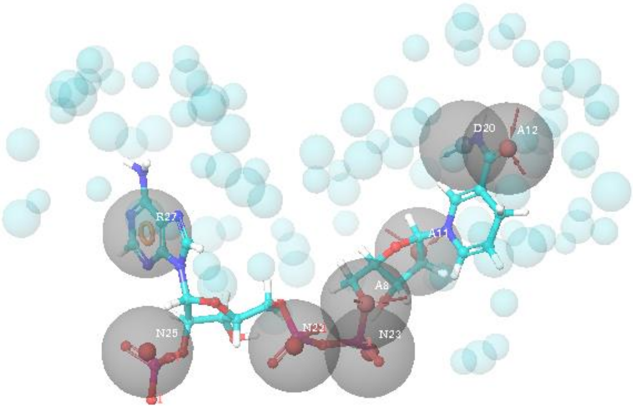

**N-Type Model1**
**D-NDP-P (Cluster 1)**

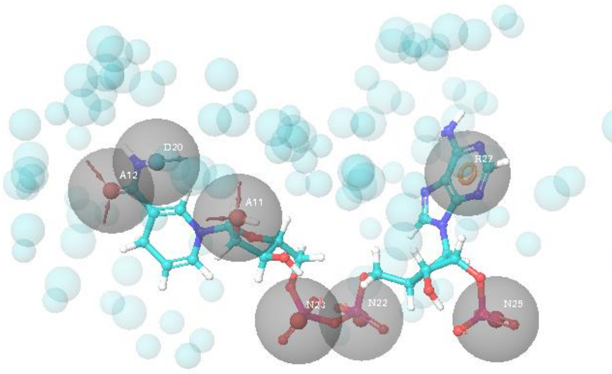

**N-Type Model2**
**D-NDP-P (Cluster 2)**

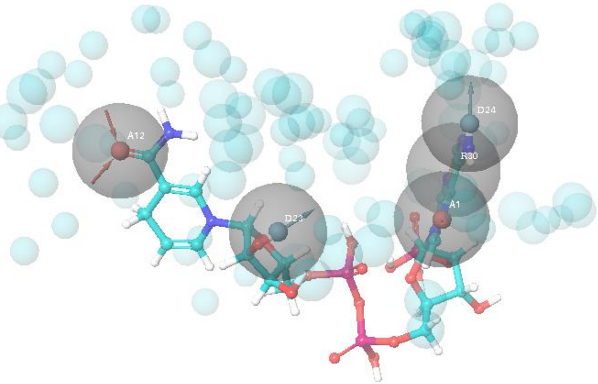

**N-Type Model3**
**D-NDP (Cluster 2)**

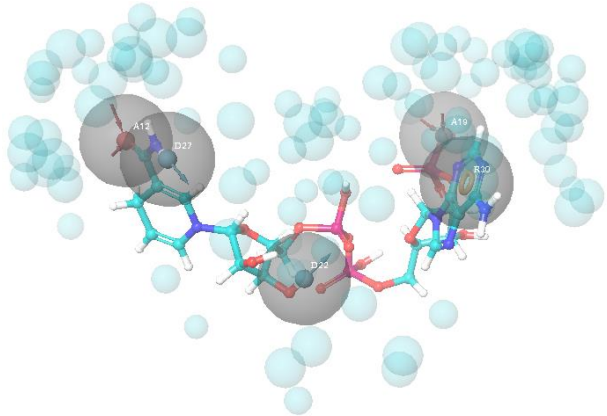

**N-Type Model4**
**D-NDP (Cluster 4)**

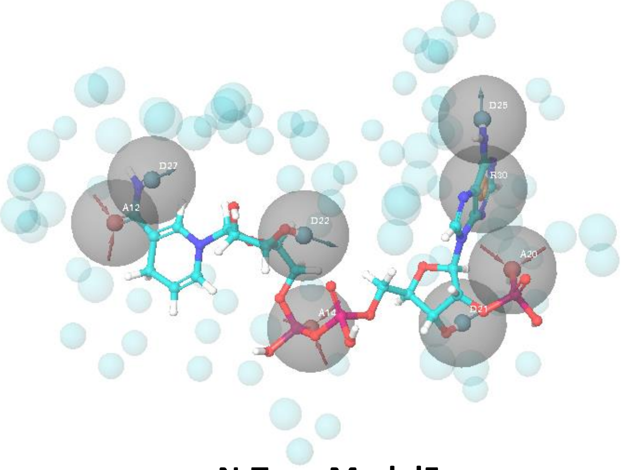

**N-Type Model5**
**D-NDP (Cluster 6)**

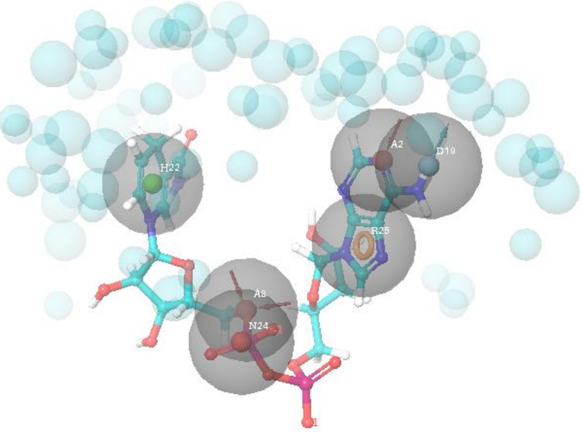

**N-Type Model6**
**D-NAI-P (Cluster 1)**

**Figure S3.** Snapshots of N-type pharmacophores with the cofactors aligned to them.

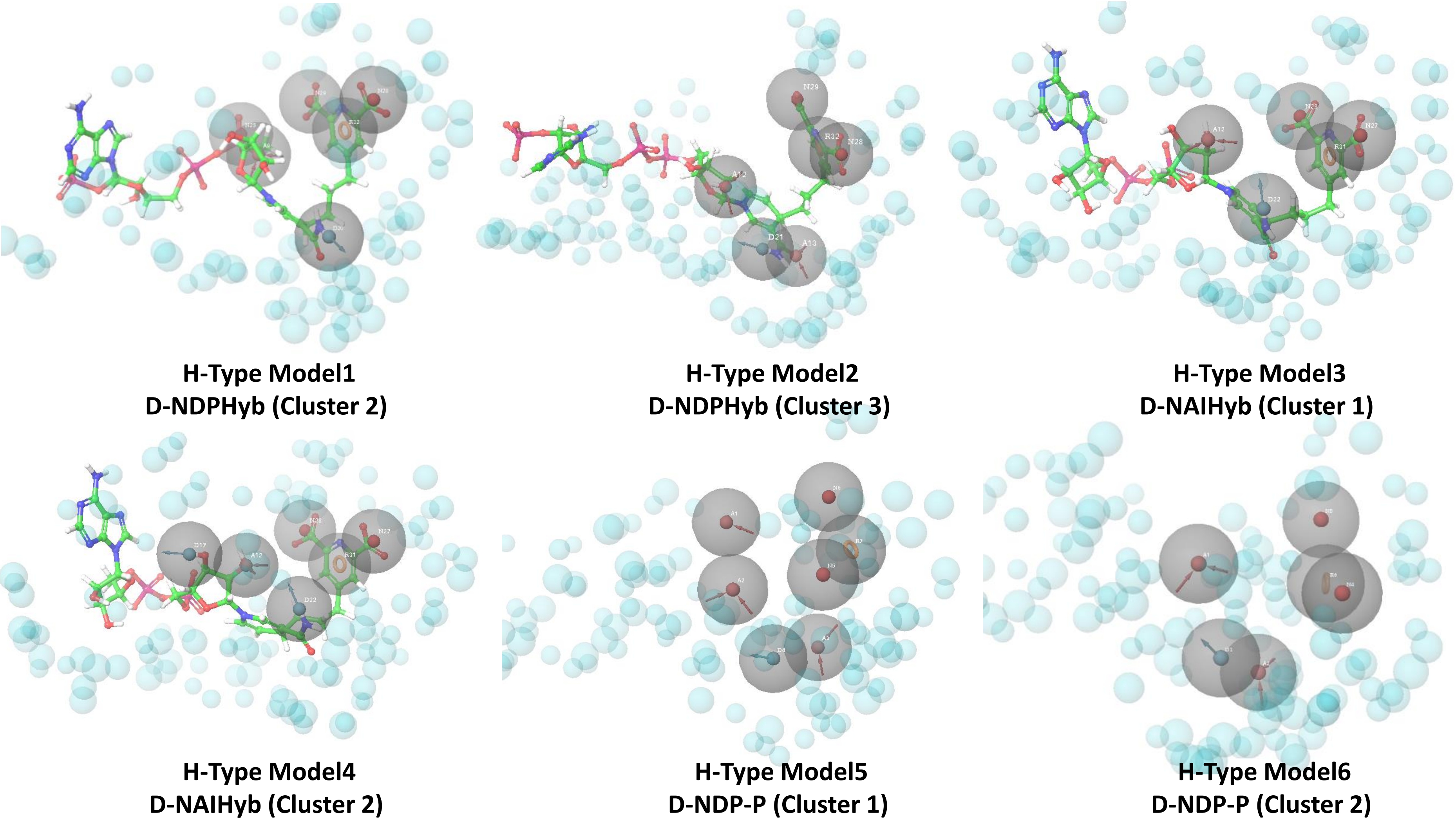


**Figure S4.** Snapshots of H-type pharmacophores with the first four aligned to the respective hybrid molecules. The last two are designed by combining two models, hence do not have a reference molecule.

**Table S3** Screening results of Mtb-active and decoy molecules

| Model | Total active molecules in the dataset | # of active molecules screened | Range of fitness score of the top screened active molecules | Total decoy molecules in the dataset | # of decoy molecules screened | Range of fitness score of the top screened decoy molecules | ROC |
|---|---|---|---|---|---|---|---|
| **H-type model** | | | | | | | |
| 1 | 2349 | 225 | 0.59 to 0.71 | 12934 | 389 | 0.44 to 0.55 | 0.10 |
| 2 | 2349 | 815 | 0.63 to 0.71 | 12934 | 1205 | 0.31 to 0.65 | 0.35 |
| 3 | 2349 | 133 | 0.64 to 0.75 | 12934 | 236 | 0.40 to 0.58 | 0.06 |
| 4 | 2349 | 286 | 0.59 to 0.66 | 12934 | 505 | 0.26 to 0.54 | 0.13 |
| 5 | 2349 | 225 | 0.51 to 0.71 | 12934 | 372 | 0.22 to 0.53 | 0.10 |
| 6 | 2349 | 874 | 0.77 to 0.83 | 12934 | 1278 | 0.29 to 0.72 | 0.37 |
| **N-type model** | | | | | | | |
| 1 | 2349 | 22 | 0.39 to 0.79 | 12934 | 31 | 0.18 to 0.53 | 0.01 |
| 2 | 2349 | 39 | 0.33 to 0.76 | 12934 | 39 | 0.2 to 0.49 | 0.02 |
| 3 | 2349 | 13 | 0.38 to 0.72 | 12934 | 14 | 0.065 to 0.61 | 0.005 |
| 4 | 2349 | 1 | 0.37 | 12934 | 2 | 0.22 to 0.44 | 0 |
| 5 | 2349 | 1 | 0.33 | 12934 | 2 | 0.28 to 0.44 | 0 |
| 6 | 2349 | 22 | 0.26 to 0.63 | 12934 | 24 | 0.01 to 0.58 | 0.01 |

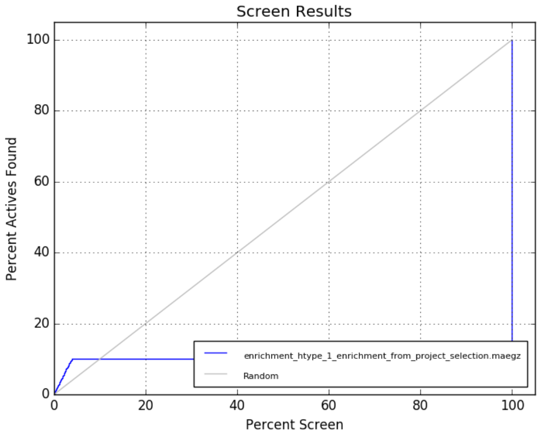


**H-type model 1**

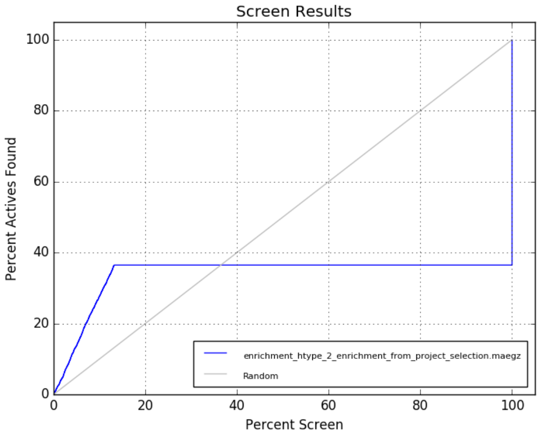


**H-type model 2**

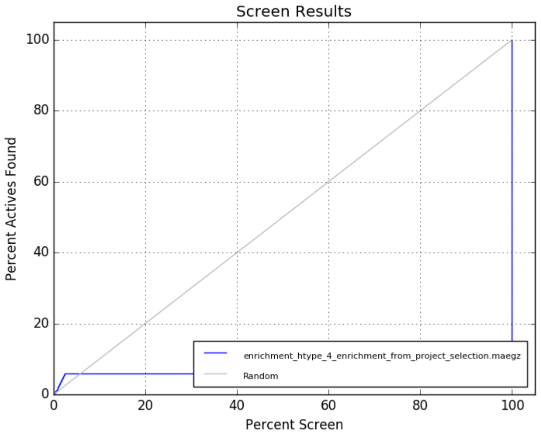


**H-type model 3**

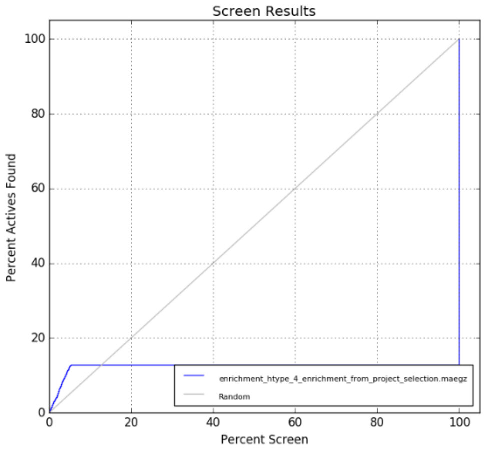


**H-type model 4**

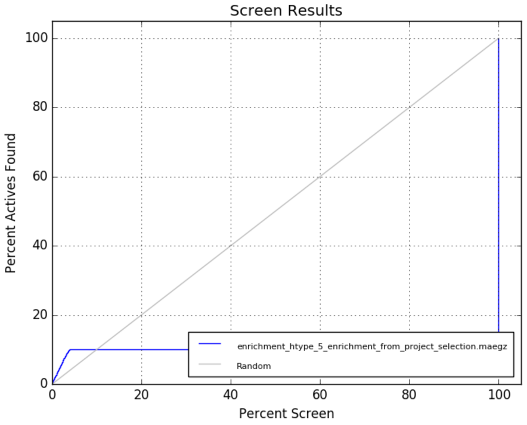


**H-type model 5**

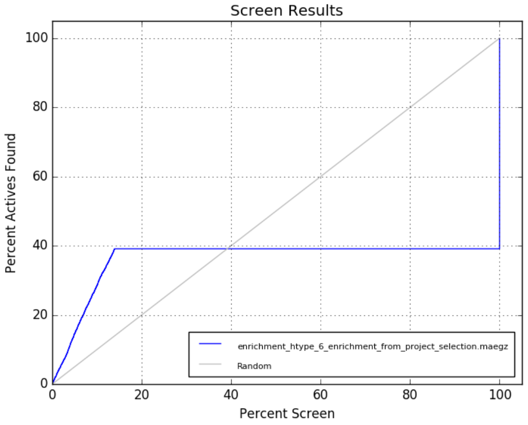


**H-type model 6**

**Figure S5.** ROC curves for Mtb-active and decoy molecules screened again H-type pharmacophore models

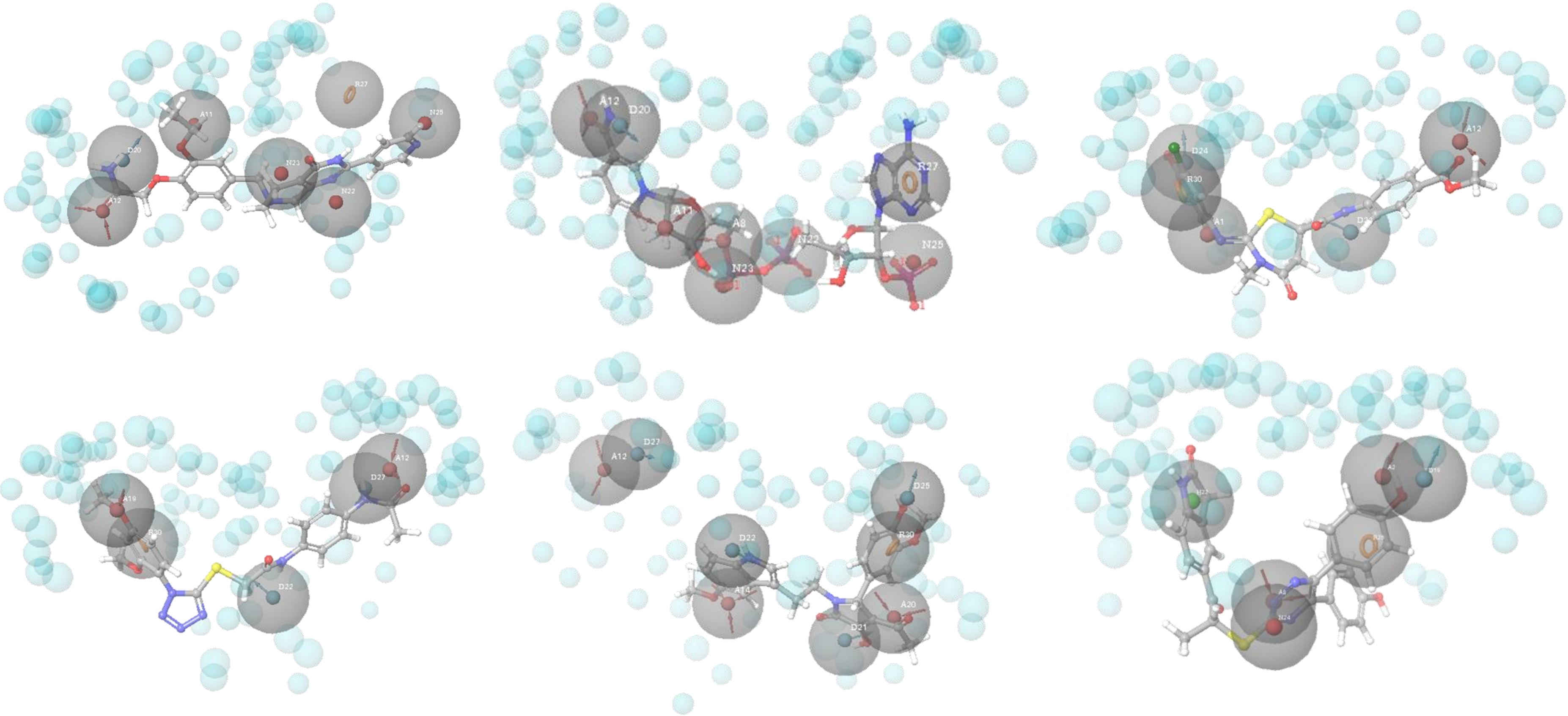

**Figure S6.** N-type pharmacophore models mapped to the respective top scoring compounds.

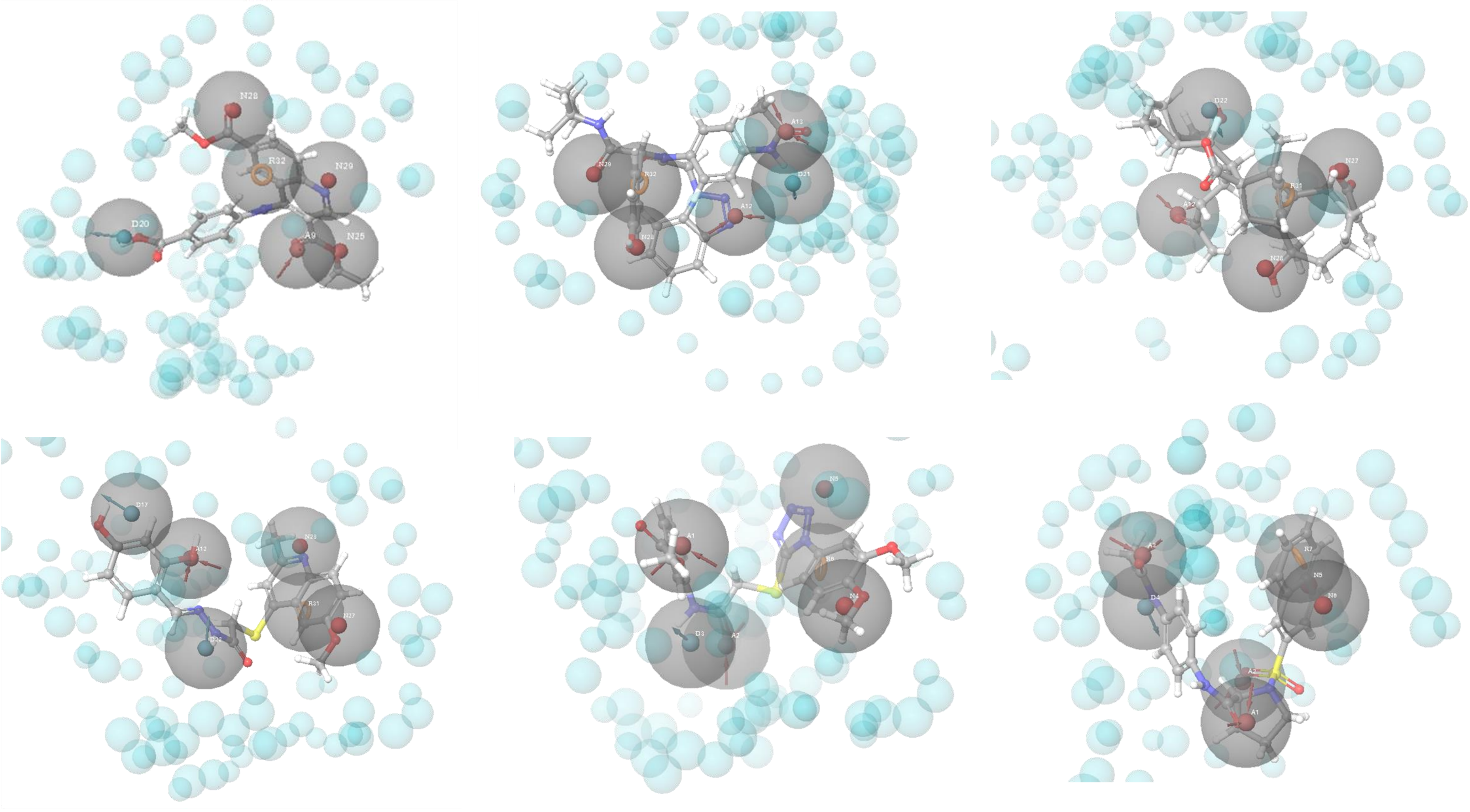


**Figure S7.** H-type pharmacophore models mapped to the respective top scoring compounds.

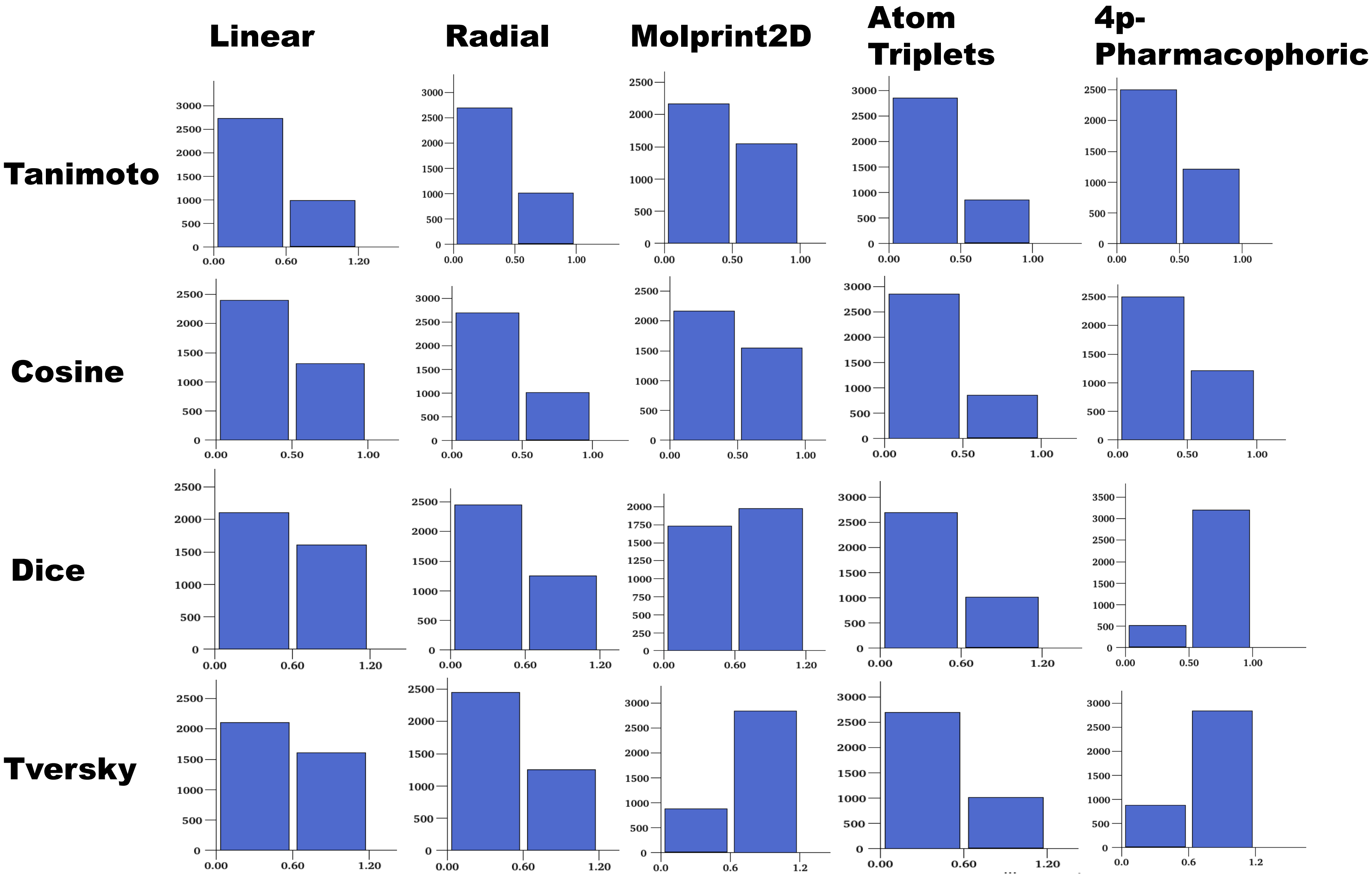


**Figure S8.** The similarity score distributions between H-set vs. N-set

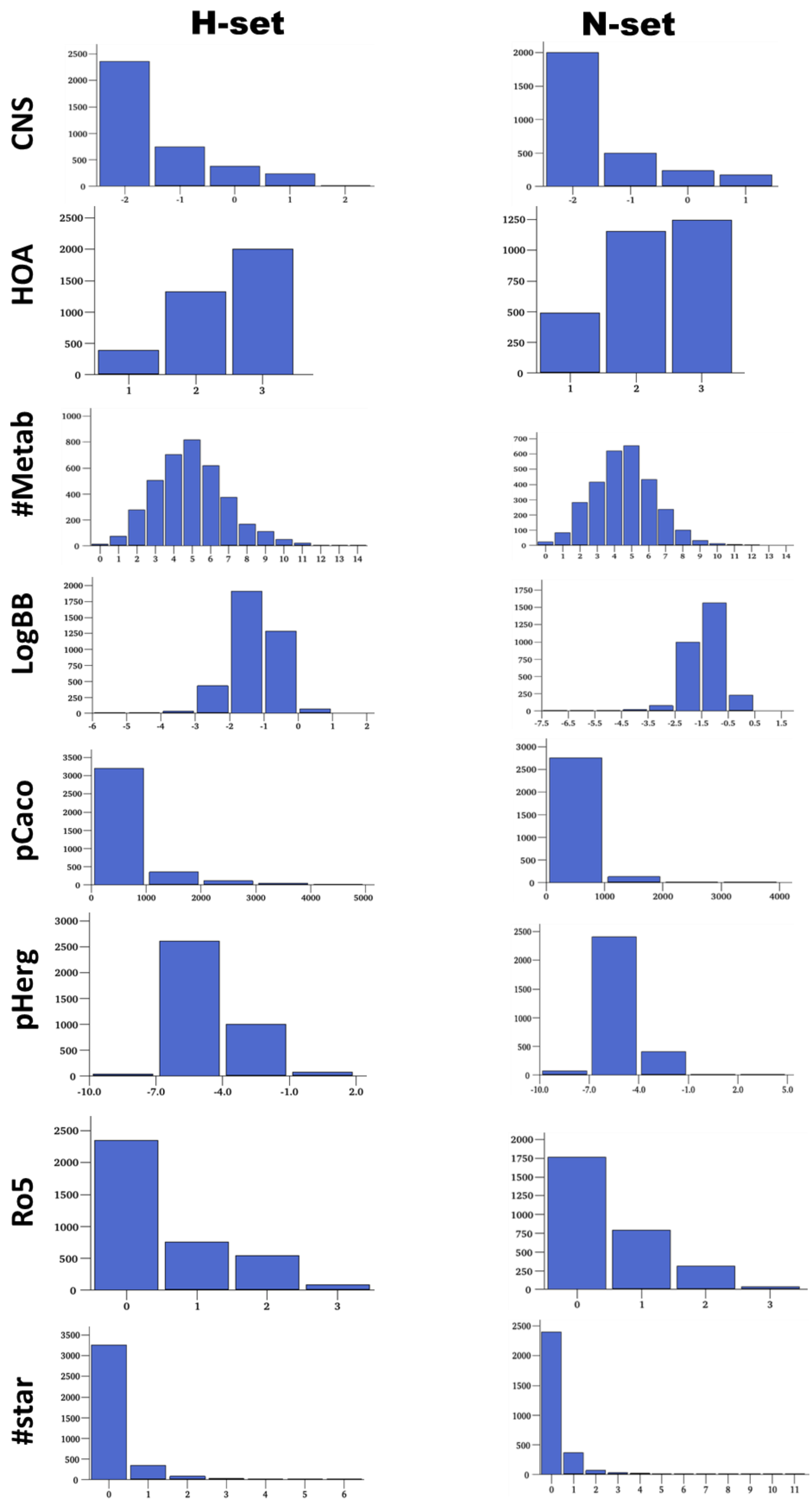


**Figure S9**. Distribution of different pharmacokinetic properties of the H-set and N-set compounds.

**List S1.** Details of all types QuikProp descriptors calculated in this study (recommended range of value for each property is mentioned in brackets).

• #stars Number of property or descriptor values that fall outside the 95% range of similar values for known drugs. The following properties and descriptors are included in the determination of #stars: MW, dipole, IP, EA, SASA, FOSA, FISA, PISA, WPSA, PSA, volume, #rotor, donorHB, accptHB, glob, QPpolrz, QPlogPC16, QPlogPoct, QPlogPw, QPlogPo/w, logS, QPLogKhsa, QPlogBB, #metabol (0 – 5)

• #rotor Number of non-trivial (not CX3), non-hindered (not alkene, amide, small ring) rotatable bonds. (0 – 15)

• mol_MW Molecular weight of the molecule. (130.0 – 725.0)

• SASA Total solvent accessible surface area (SASA) in square angstroms using a probe with a 1.4 Å radius. (300.0 – 1000.0)

• FOSA Hydrophobic component of the SASA (saturated carbon and attached hydrogen). (0.0 – 750.0)

• FISA Hydrophilic component of the SASA (SASA on N, O, and H on heteroatoms). (7.0 – 330.0)

• PISA (carbon and attached hydrogen) component of the SASA. (0.0 – 450.0)

• WPSA Weakly polar component of the SASA (halogens, P, and S). (0.0 – 175.0)

• volume Total solvent-accessible volume in cubic angstroms using a probe with a 1.4 Å radius. (500.0 – 2000.0)

• donorHB Estimated number of hydrogen bonds that would be donated by the solute to water molecules in an aqueous solution. Values are averages taken over a number of configurations, so they can be non-integer. (0.0 – 6.0)

• accptHB Estimated number of hydrogen bonds that would be accepted by the solute from water molecules in an aqueous solution. Values are averages taken over a number of configurations, sothey can be non-integer. (2.0 – 20.0)

• glob Globularity descriptor. (0.75 – 0.95)

• QPpolrz Predicted polarizability in cubic angstroms. (13.0 – 70.0)

• QPlogPC16 Predicted hexadecane/gas partition coefficient. (4.0 – 18.0)

• QPlogPoct‡ Predicted octanol/gas partition coefficient. (8.0 – 35.0)

• QPlogPw Predicted water/gas partition coefficient. (4.0 – 45.0)

• QPlogPo/w Predicted octanol/water partition coefficient. (–2.0 – 6.5)

• QPlogS Predicted aqueous solubility, log S. S in mol dm–3 is the concentration of the solute in a saturated solution that is in equilibrium with the crystalline solid. (–6.5 – 0.5)

• CIQPlogS Conformation-independent predicted aqueous solubility, log S. S in mol dm–3 is the concentration of the solute in a saturated solution that is in equilibrium with the crystalline solid. –6.5 – 0.5

• QPlogHERG Predicted IC50 value for blockage of HERG K+ channels. (concern below –5)

• QPPCaco Predicted apparent Caco-2 cell permeability in nm/sec. Caco-2 cells are a model for the gut-blood barrier. QikProp predictions are for non-active transport. (<25 poor, >500 great)

• QPlogBB Predicted brain/blood partition coefficient. ( –3.0 – 1.2)

• QPPMDCK Predicted apparent MDCK cell permeability in nm/sec. MDCK cells are considered to be a good mimic for the bloodbrain barrier. QikProp predictions are for non-active transport. (<25 poor, >500 great)

• QPlogKp Predicted skin permeability, log Kp. ( –8.0 – 1.0)

• QPlogKhsa Prediction of binding to human serum albumin. (–1.5 – 1.5)

• HumanOralAbsorption Predicted qualitative human oral absorption: 1, 2, or 3 for low, medium, or high.

• PercentHuman- OralAbsorption Predicted human oral absorption on 0 to 100% scale. The prediction is based on a quantitative multiple linear regression model. This property usually correlates well with HumanOralAbsorption, as both measure the same property. ( >80% is high ,<25% is poor)

• PSA Van der Waals surface area of polar nitrogen and oxygen atoms. (7.0 – 200.0)

• RuleOfFive Number of violations of Lipinski's rule of five. The rules are: mol_MW < 500, QPlogPo/w < 5, donorHB ≤ accptHB ≤10. Compounds that satisfy these rules are considered druglike. (maximum is 4)
